# Perturbing nuclear glycosylation in the mouse preimplantation embryo slows down embryonic growth

**DOI:** 10.1101/2024.01.22.576677

**Authors:** Sara Formichetti, Urvashi Chitnavis, Agnieszka Sadowska, Na Liu, Ana Boskovic, Matthieu Boulard

**Affiliations:** Epigenetics and Neurobiology Unit, European Molecular Biology Laboratory (EMBL), Monterotondo, Italy; Collaboration for joint PhD degree between EMBL and Heidelberg University, Germany

**Author notes:** Correspondence (A.B.), (M.B.).

## Abstract

The only known form of intracellular protein glycosylation (O-GlcNAc) is reversible and has been mapped on thousands of cytoplasmic and nuclear proteins, including RNA polymerase II, transcription factors and chromatin modifiers. The O-GlcNAc modification is catalyzed by a single enzyme known as O-GlcNAc Transferase (OGT), that is required for mammalian early development. Remarkably, the regulatory function of protein O-GlcNAcylation in the embryo as well as the embryonic O-GlcNAc proteome remain unknown. Here, we devised a new method to enzymatically remove O-GlcNAc from preimplantation embryonic nuclei, where it accumulates coincidently with embryonic genome activation (EGA). Unexpectedly, the depletion of nuclear O-GlcNAc to undetectable levels has no impact on EGA, but dampens the transcriptional activation of the translational machinery, and triggers a spindle checkpoint response. These molecular alterations were phenotypically associated with a developmental delay starting from early cleavage stages and persisting after embryo implantation, establishing a novel link between nuclear glycosylation and embryonic growth.

## Introduction

Following fertilization, the embryo relies on the payload of RNA and proteins carried by the oocyte. Then, it undergoes genome-wide reprogramming involving changes to chromatin organization as well as reconfiguration of the transcriptome, with the degradation of maternal transcripts and the activation of the embryonic genome. A less studied concomitant phenomenon is the reconfiguration of the proteome, which includes dynamic post-translational modifications (PTMs). Among PTMs, intracellular glycosylation remains uncharacterized in the early embryo.

The main form of intracellular glycosylation in animals is O-GlcNAcylation, the reversible addition of the monosaccharide O-linked N-acetylglucosamine (O-GlcNAc) to specific threonine and serine residues. A single pair of enzymes catalyzes the deposition and removal of O-GlcNAc, namely O-GlcNAc transferase (OGT)^1,2^ and O-GlcNAc hydrolase (OGA)^3^, both encoded by one single gene with the only known exception of zebrafish^4^. In Drosophila, loss-of-function of *Ogt* (also known as *super sex combs* in fly genetics) produces a stereotypical post-gastrulation developmental phenotype characterized by the expression of homeotic genes outside their normal boundaries^5^. In mammals, *Ogt* knockout causes a more severe phenotype: the maternal inheritance of a non-functional copy of *Ogt* leads to embryonic lethality around the blastocyst stage^6^, implying that mammalian OGT and/or O-GlcNAc are essential either for oocyte development or for preimplantation processes. However, the early lethal phenotype has obstructed the understanding of O-GlcNAc function in the mammalian embryo, where the relevance of this PTM for the different cellular and developmental processes remains unknown.

O-GlcNAc is a highly pleiotropic modification, mapped on thousands of mammalian proteins^7^ and involved in many pathways, including translation^8–10^, glucose catabolism^11–13^ and the cell cycle^14^. In the nucleus, O-GlcNAc modifies numerous transcription factors, among which the developmentally relevant OCT4 and SOX2 are noteworthy^15^. OCT4 is required for the maintenance of pluripotency in the inner cell mass (ICM) of the blastocyst^16^; SOX2 is one of the first transcription factors to function asymmetrically, already at the 4-cell stage, in those cells that will give rise to the ICM^17^. Because O-GlcNAcylation of OCT4 has been shown to be required for its reprogramming function *in vitro*^15^, O-GlcNAc might contribute to the establishment of the pluripotency gene network during preimplantation development. Another noticeable nuclear target of OGT is the C-terminal domain (CTD) of RNA Polymerase II, where O-GlcNAc was mapped on threonine 4 and serine 5^18^. In spite of its presence of key gene expression regulators, it remains unclear whether O-GlcNAc plays an instructive function in controlling transcriptional activity^19^. The preimplantation arrest resulting from *Ogt* loss-of-function is consistent with an hypothetical role of O-GlcNAc in the first transcriptional event of the embryo, namely the embryonic genome activation (EGA), for which all players are still unknown in mammals^20^.

Finally, the donor substrate for O-GlcNAc is UDP-GlcNAc, the end product of the hexosamine biosynthetic pathway, which channels 2-3% of intracellular glucose and whose flux is responsive to levels of nutrients such as glucose and glutamine^21,22^. For this reason, O-GlcNAc has been recurrently associated with nutrient sensing and cellular adaptation^11,12^. The early mammalian embryo is a biological paragon of a highly metabolically dynamic system. Glucose import is low until the 8-cell stage and glycolysis almost undetectable during preimplantation development, when the embryo mostly relies on ATP production through oxidative phosphorylation^23^. Accordingly, the expression of enzymes participating in oxidative phosphorylation is upregulated at EGA and gradually increases during the following cleavages^24^. At implantation, the decrease in oxygen level together with the need for high proliferation rate and biomass production is accompanied by a metabolic switch which favors glycolysis^24^. In light of these findings, early development has the potential to be regulated by a metabolically-sensitive PTM such as O-GlcNAc.

The above hypotheses on the possible roles of mammalian O-GlcNAc in early development are based on *in vitro* evidence but have not been tested *in vivo*. This is largely due to the hurdle posed to classical genetics approaches by the requirement of the *Ogt* maternal allele. In this study, we documented for the first time the dynamics of O-GlcNAc across key preimplantation stages, which highlighted its enrichment in the nucleus from the 2-cell stage onwards. We then developed a strategy to enzymatically remove both embryonic and maternally inherited O-GlcNAc from the nuclei of the growing preimplantation embryo, and studied the resulting molecular and morphological phenotype at key preimplantation stages and up to postimplantation. Importantly, our approach targets virtually the entire nuclear O-GlcNAc proteome, addressing the pleiotropy of this modification. Unexpectedly, our findings revealed that disturbing O-GlcNAc homeostasis does not affect EGA, nor the developmental gene networks, but impairs the transcriptional activation of the translational machinery and slows down embryonic growth, starting from cleavage stages and persisting after embryo implantation.

## Results

### Uncoupled dynamics of OGT and O-GlcNAc across mouse preimplantation development

The dynamics of protein O-GlcNAcylation in the early mouse embryo is unknown. Thus, we set out to determine the spatiotemporal profile of OGT (Figure 1A) and O-GlcNAc (Figure 1B) across key preimplantation stages. Embryos were generated through in vitro fertilization (IVF) and the following stages were collected for immunofluorescence staining: zygotes at different pronuclear (PN) stages (0-4 h post-IVF), late 2-cell embryos (22-26 h post-IVF), morulae (48 h post-IVF) and late blastocysts (68 h post-IVF). We also collected unfertilized MII oocytes for analysis.

**Figure 1.**
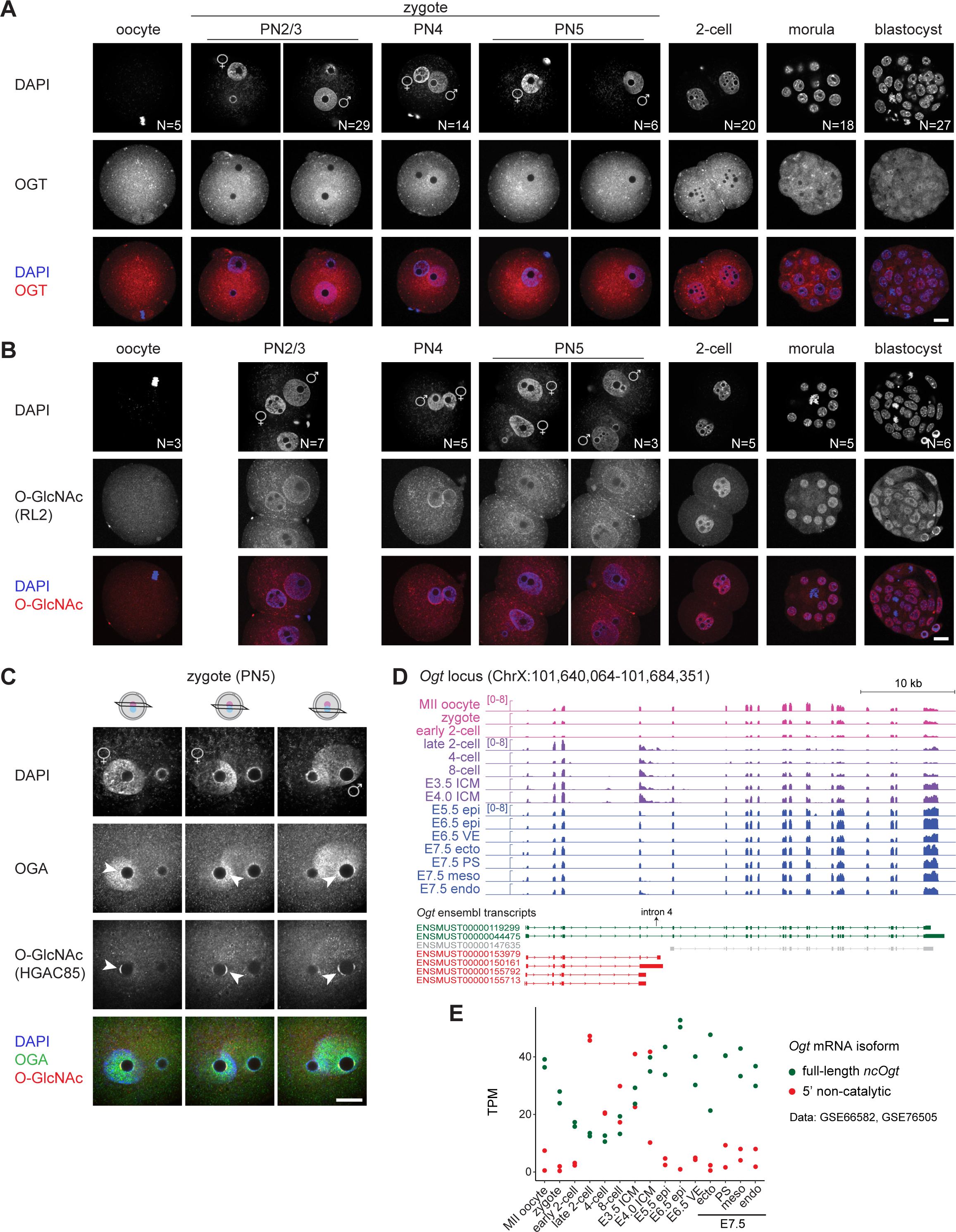
Uncoupled dynamics of OGT and O-GlcNAc across mouse preimplantation development. (A,B) Immunofluorescence staining of OGT (ab177941) (A) and the O-GlcNAc modification (RL2 antibody) (B) in MII oocytes, zygotes at various pronuclear stages, 2-cell embryos (22-26 h post-IVF), morulae (48 h post-IVF) and blastocysts (68 h post-IVF) generated through IVF. One z-plane is shown for each embryo, except for some zygotes for which two z-planes are shown. (C) Immunofluorescence co-staining of O-GlcNAc (HGAC85 antibody) and OGA. Co-localization of the two signals at perinucleolar foci in both parental pronuclei is indicated by white arrows. Different z-planes are shown for the same zygote; note that the signal is visible only on certain nuclear planes. Five zygotes were imaged, with the same result. (A-C) Embryos were mounted on coverslips and imaged using a scanning confocal microscope. Scale bar indicates 20 μm. PN = pronuclear stage. DNA was stained with DAPI. The total number of embryos imaged is indicated for each stage and comes from at least two independent IVF experiments. (D) Genome browser view of the *Ogt* gene showing the alignment of mRNA-Seq reads from GSE6658232 (MII oocyte to E4.0 ICM) and GSE76505^33^ (E5.5 to E7.5). Read counts are normalized to bins per million. One biological replicate per stage is shown. (E) Salmon quantification of the Ogt transcript isoforms from the same datasets as in (D). TPM of isoforms ENSMUST00000119299 and ENSMUST00000044475 were averaged (full-length ncOgt), as well as TPM of ENSMUST00000153979, ENSMUST00000150161, ENSMUST00000155792 and ENS-MUST00000155713 (5’ non-catalytic). ENSMUST00000147635 was excluded from the plot because of no evidence of its presence based on a manual inspection of the IGV tracks. (D,E) epi = epiblast, ecto = ectoderm, PS = primitive streak, meso = mesoderm, endo = endoderm.

Firstly, we observed a striking enrichment of the OGT signal in the paternal pronucleus of the zygote compared to the maternal one at all PN stages, consistent with a previous study reporting this asymmetry at PN3/4^25^. The same pattern was found with embryos produced by natural mating (Figure S1A). In contrast, the O-GlcNAc signal was equivalently present on both pronuclei, as was the signal of the O-GlcNAc hydrolase OGA (Figure S1B), pointing to a potential catalytically-independent role of OGT in the zygote. Secondly, the nuclear-to-cytosolic signal ratio of O-GlcNAc markedly increases from the 2-cell stage onwards, encouraging a further investigation of the role of this modification in the embryonic nuclei specifically. The OGT signal is instead roughly equivalent between the nucleus and the cytoplasm from the 2-cell stage onwards, while OGA is more abundant in the nucleus (Figure S1B). It is noteworthy that OGT, OGA and O-GlcNAc were all undetectable on the metaphase chromosomes, while the OGT but not the OGA signal was enriched on the oocyte meiotic spindle (oocytes in Figures 1A and S1B), as previously reported^26^.

In the zygote, O-GlcNAc displayed an intense signal at the nuclear membrane. This is likely attributable to the highly O-GlcNAcylated state of the FG nucleoporins (FG-Nups) subunits of the nuclear pore complex (NPC)^27,28^, since the antibody anti-O-GlcNAc RL2 has been raised against rat NPC-lamina nuclear fraction^29^. However, zygotic nuclear O-GlcNAc was not found exclusively at the NPC, as shown by a different antibody (clone HGAC85) that stained perinucleolar foci in the zygote and in the 2-cell embryo (Figures 1C and S1C). The perinucleolar signal was lost after enzymatic removal of O-GlcNAc (outlined below) (Figure S1C), proving the specificity of this O-GlcNAc pattern. Remarkably, perinucleolar O-GlcNAc overlapped with the OGA signal (Figure 1C), suggesting an active remodeling of nucleoli-associated clusters of O-GlcNAcylated proteins in the zygote.

The monoclonal antibody against OGT used in this study recognizes the C-terminus of OGT, thus it should detect all reported OGT isoforms, which include the nuclear-cytoplasmic one (ncOGT) in addition to a mitochondrial isoform and a shorter isoform of unknown localization and function^30,31^. To gain insight into OGT isoform usage in the early embryo, we analyzed publicly available mRNA-Seq data spanning mouse pre- and post-implantation stages^32,33^ (Figure 1D). We noted that the longer transcript isoforms (in green in Figure 1D), which encode ncOGT, are the ones predominantly present in the oocyte, after which their levels gradually diminish and start to rise again only from the 8-cell stage (Figure 1E). Instead, a shorter form of *Ogt* terminating after exon 4 is produced starting from EGA and is present at all cleavage stages (Figures 1D, 1E and S1D). This transcript isoform could correspond to the annotated shorter N-terminal ones annotated in Ensembl (in red in Figure 1D), which are predicted to be non-coding or catalytically inactive. Of note, this shorter *Ogt* transcript retains intron 4 of the full-length coding isoform, a phenomenon previously shown to increase with higher O-GlcNAc levels and associated with nuclear degradation of the transcript, in turn maintaining O-GlcNAc homeostasis^34^.

The same analysis for the *Oga* locus showed that during preimplantation stages two mRNA isoforms are produced, one not including exon 11 as well as the C-terminal exons of the longest isoform (Figures S1E and S1F). The inclusion of *Oga* exon 11 has been reported to correlate with functional *Oga* splicing and, notably, to increase in conditions of higher O-GlcNAc, as opposed to the inclusion of *Ogt* intron 4^35^.

In summary, the subcellular pattern of the O-GlcNAc modification is highly dynamic during mouse preimplantation development, and distinct from that of OGT. The concentration of the latter appears to be tightly controlled in the early embryo, first by its confinement in the paternal pronucleus and then through alternative production of RNA isoforms.

### Effective enzymatic depletion of O-GlcNAc from mouse embryonic nuclei

The high abundance of O-GlcNAc-modified proteins in the nucleus from the 2-cell stage onwards (Figure 1B) prompted us to functionally assess the significance of nuclear O-GlcNAcylation during preimplantation development. To this aim, we needed an experimental system that overcomes embryonic lethality of maternal *Ogt* loss-of-function and eliminates the payload of protein O-GlcNAcylation inherited from the oocyte.

Our strategy was to enzymatically remove the O-GlcNAc moieties from the zygote onwards by overexpressing a recombinant O-GlcNAcase targeted to the nucleus. We leveraged the homologous O-GlcNAcase from the human gut symbiont *Bacteroides thetaiotaomicron* (BtGH84), whose structure and catalytic mechanism have been well-characterized *in vitro*^36^ and which has been shown to remove O-GlcNAc at targeted mammalian genomic loci when fused to dCas9^37^. We fused BtGH84 to N-terminal and C-terminal nuclear localization signals (NLS) and to an EGFP reporter at the N-terminus and injected the mRNA of this construct (henceforth called “Btgh”) into zygotes 2 hours after IVF (∼PN3; Figure 2A). Two other groups of embryos were used to control for the effects due to the injection procedure and the translation of an exogenous protein: non-injected embryos and embryos injected with a catalytically-inactive Btgh construct bearing a single amino acid mutation (D242A)^36^ (henceforth “dBtgh” for “dead Btgh”). After injections of Btgh, the O-GlcNAc signal was dramatically reduced to undetectable levels already at the early 2-cell stage (i.e. before EGA; Figures 2B and 2C), while it remained comparable to non-injected embryos following dBtgh expression. We confirmed the O-GlcNAc depletion to undetectable levels using a different antibody against O-GlcNAc (clone HGAC85; Figure S1C). Importantly, O-GlcNAc depletion persisted until the late blastocysts stage (embryonic day 4; E4), the last stage assessed with immunostaining (Figure 2D).

**Figure 2.**
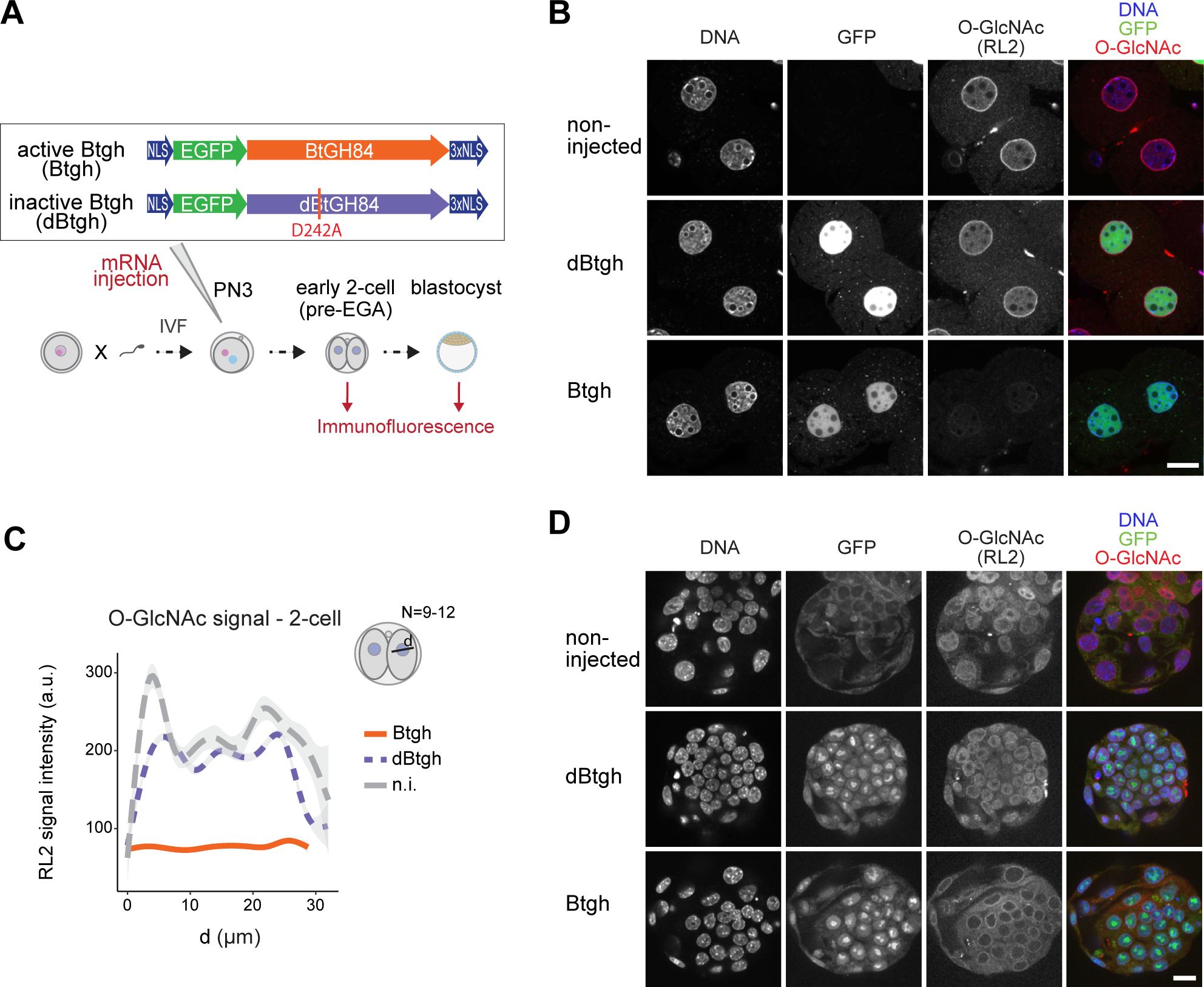
Effective enzymatic depletion of O-GlcNAc from mouse embryonic nuclei. (A) Experimental design for results in (B-D). Zygotes were generated through IVF and injected 2 h later with the mRNA of the active or inactive form of NLS-EGFP-BtGH84-3xNLS (Btgh and dBtgh, respectively), or left uninjected. The three groups of embryos were cultured ex-vivo and fixed either as pre-EGA 2-cell or at the blastocyst stage (68 h post-IVF). (B,C) Confocal imaging of pre-EGA 2-cell embryos (B) and blastocysts (D) from zygotes injected with Btgh/dBtgh or non-injected, stained with an anti-O-GlcNAc antibody (RL2). Embryos were mounted on coverslips and imaged using a scanning confocal microscope. One z-plane is shown for each embryo. Scale bar indicates 20 μm. DNA was stained with DAPI. The GFP signal comes from the Btgh/dBtgh protein fused to GFP. (C) Quantification of the O-GlcNAc signal across single nuclei of 2-cell embryos from the experiment in (B). N = 9-12 nuclei per experimental group were quantified, from two independent microinjection experiments.

### Nuclear O-GlcNAc depletion does not affect differentiation but slows down development

Having validated the efficiency of our novel strategy, we investigated the effect of nuclear O-GlcNAc depletion on the progression of preimplantation development. We found no impact on blastocyst formation rate *ex vivo* at E4 (Figure S2A), nor on the specification of the trophectoderm (TE) and inner cell mass (ICM) lineages (Figure S2B). Hence, physiological levels of nuclear O-GlcNAc are dispensable for progression to the blastocyst stage. This result was unexpected because glycosylation of the master regulator of pluripotency OCT4 was shown to be required for its reprogramming function *in vitro*^15^.

Next, since *Ogt*-null embryos die around the time of implantation (E4-5)^6^, we asked whether nuclear O-GlcNAc might be vital for peri-implantation processes. Zygotes injected with either the Btgh or dBtgh mRNA were grown overnight, and the early 2-cell embryos surgically transferred into pseudopregnant females and dissected six days later (embryonic day 7; Figure 3A). The percentage of embryos recovered at E7 between the two groups of injections was not significantly different (Btgh: 43%, N=92, dBtgh: 48%, N=134; p-value = 0.64; Figure 3B), indicating that the depletion of nuclear O-GlcNAc had no impact on implantation. Although staging of the embryos based on the appearance of the mesoderm^38^ showed a similar percentage of embryos to be Pre-, Early-or Mid-Streak stages in both groups (Table S1), nuclear O-GlcNAc-depleted E7 embryos were on average smaller (Figure 3C). This difference was mainly driven by 2 out of 5 litters of Btgh-injected embryos, thus the phenotype might be partially penetrant.

**Figure 3.**
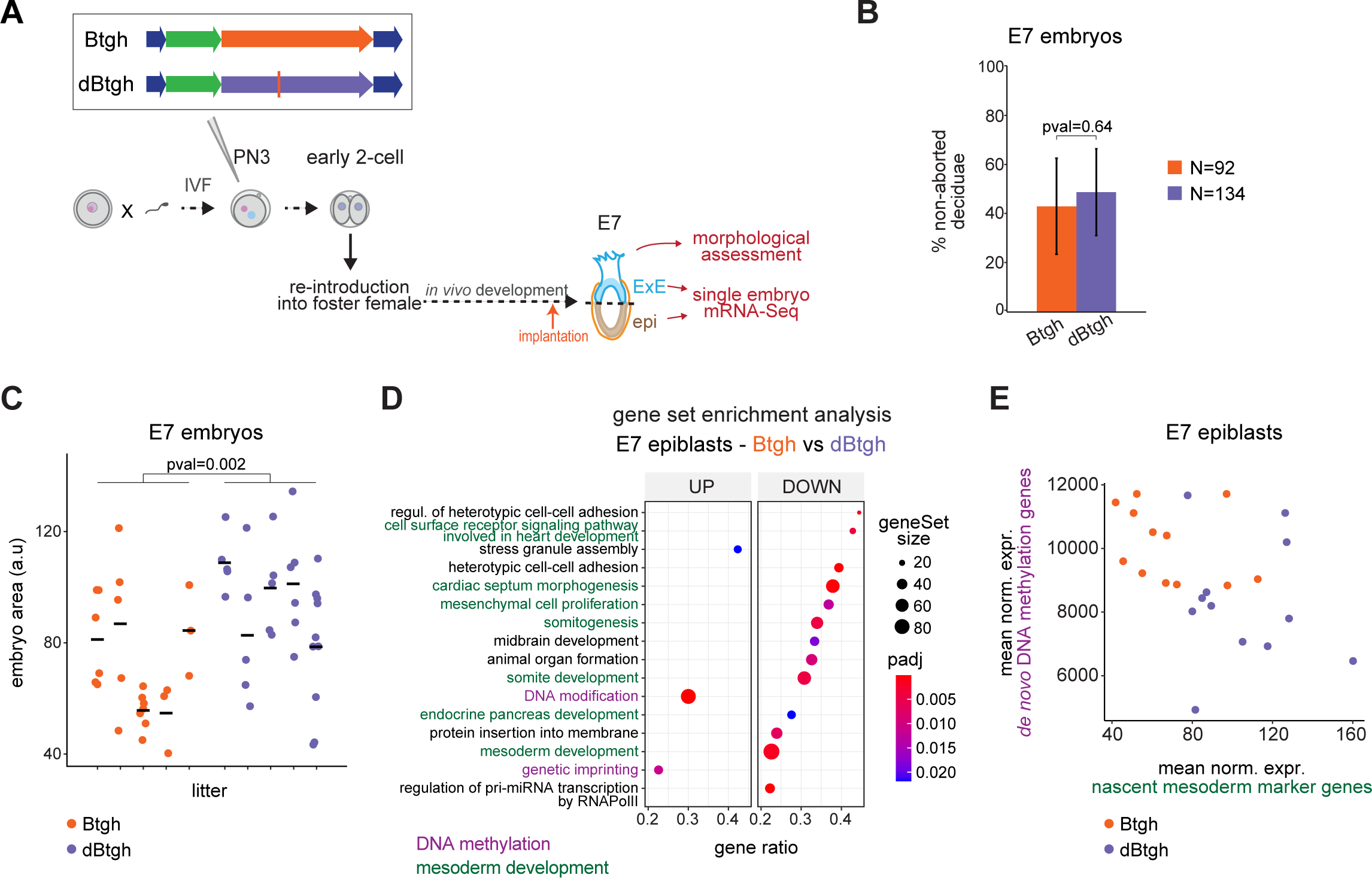
Nuclear O-GlcNAc depletion does not affect differentiation but slows down development. (A) Experimental design for results in (B-F). Zygotes were generated through IVF and injected 2 h later with the mRNA of either Btgh or dBtgh, then cultured ex-vivo overnight and the next morning transferred to pseudopregnant females. Six days later (seven days after IVF, thus embryonic day 7) the embryos were dissected, imaged and cut into two halves, largely corresponding to the epiblast (epi) and extraembryonic ectoderm (ExE) tissues, respectively. The two halves were separately processed for single-embryo mRNA-Seq. (B) Percentage of Btgh/dBtgh-injected embryos recovered at E7. N = 92 and 134 total injected 2-cell embryos transferred to pseudopregnant females for Btgh and dBtgh, respectively. Bar heights indicate the average, and error bars indicate the standard deviation of five replicates of the microinjection experiment. P-value was computed using unpaired Student’s t-test, assuming unequal variance. (C) Size of the Btgh/dBtgh-injected E7 embryos for all dissected litters, as measured by the area of the embryos in the images taken at the dissection microscope (Figure S2C, upper part). Each column shows a litter. P-value was computed using unpaired Wilcoxon rank sum exact test. (D) Gene set enrichment analysis of gene expression changes in E7 epiblasts derived from Btgh-injected embryos versus dBtgh-injected controls. Among the significant Biological Process gene ontology (GO) terms, the 16 with the highest Normalized Enrichment Score are shown, ordered by gene ratio and colored based on manual grouping of cellular functions. Size of dots is proportional to the number of total genes of a GO term. Gene ratio = fraction of total genes of the GO term which are concordantly changing between the two conditions. (E) Mean DESeq2-normalized counts of E7 mesoderm markers from https://marionilab.cruk.cam.ac.uk/-MouseGastrulation2018/^44^ versus mean DESeq2-normalized counts of enzymes associated with de novo DNA methylation (*Dnmt3a*, *Dnmt3b*, *Tet1*) in single E7 epiblasts.

To gain molecular insights into the observed phenotype, we microdissected individual E7 conceptuses from the two experimental groups into the epiblast and extraembryonic ectoderm (ExE) halves, and processed the single halves from single embryos for mRNA-Seq (Figure S2C). Of note, we interpret gene expression differences between the two groups of postimplantation embryos as a consequence of the depletion of O-GlcNAc during preceding stages rather than as a result of active ongoing removal, because the Btgh mRNA and protein have been highly diluted by E7. Accordingly, the injected Btgh/dBtgh mRNA was no longer detectable at E7, while a low reads count could still be detected in E4 blastocyst data (Figure S2D). Moreover, a compensatory change in expression of the O-GlcNAc enzymes has been reported whenever O-GlcNAc homeostasis is disturbed^34,39–43;^ in our data, *Ogt* was found upregulated and *Oga* downregulated in Btgh-injected embryos at the morula and blastocyst stages, while this phenomenon disappeared at E7 (Figure S2E), arguing that O-GlcNAc homeostasis has been restored after implantation.

While no difference in gene expression was observed in the ExE between the two groups, several genes were differentially expressed in the epiblast (adj. p-value < 0.05, any log_2_FC; DEGs; Figure S2F). Gene set enrichment analysis (GSEA) in this tissue uncovered downregulation of genes associated with mesoderm development and upregulation of genes involved in DNA methylation upon Btgh injection (Figure 3D). The development of the mesoderm characterizes the transition between E6.5 and E7.5^44^. On the other hand, the onset of mammalian gastrulation coincides with *de novo* DNA methylation, with the enzymes involved in this process peaking in expression at E5.5 and then gradually decreasing (Figure S2G). Hence, the transcriptome of the epiblasts from Btgh-injected embryos resembles that of a slightly earlier developmental stage, implying a developmental delay. In agreement, all significantly upregulated genes are found to decrease in expression during the transition from E5.5 to E7.5 in unperturbed embryos, while the opposite is true for the downregulated DEGs (Figure S2H). We used the expression of E7 mesodermal markers^44^ and DNA methylation enzymes (*Dnmt3a*/*b*, *Tet1*) as a single-embryo measure of the developmental delay suggested by both morphological assessment and differential gene expression analysis. Nine out of twelve epiblasts isolated from Btgh-injected embryos showed lower expression of E7 mesodermal markers compared to controls (and in most cases also higher expression of *Dnmt*s and *Tet1*; Figure 3E), thus supporting a highly penetrant developmental delay after implantation upon nuclear O-GlcNAc depletion preimplantation.

### Nuclear O-GlcNAc is dispensable for EGA

As noted above, the postimplantation phenotype of O-GlcNAc-depleted embryos is a consequence of a perturbation caused by the absence of nuclear O-GlcNAc during earlier stages (Figures S2D and S2E). Hence, we assessed the developmental origin of the growth retardation observed at E7 by analyzing individual O-GlcNAc-depleted preimplantation embryos collected at the post-EGA 2-cell, morula and blastocyst stages via single-embryo mRNA-Seq (Figure 4A). Notably, principal component analysis (PCA) of all embryo transcriptomes from the three stages revealed that O-GlcNAc-depleted morulae were shifted farther from the blastocysts compared to their control counterparts (Figure 4B). The direction of PC1 and PC2 correlates with developmental progression, as supported by the dynamics of genes contributing to these PCs (Figure 4C). Therefore, this result points to a developmental delay already captured in the morulae transcriptomes. Taken together, we conclude that nuclear O-GlcNAc removal across preimplantation does not prevent cell differentiation nor implantation, but causes a slight developmental delay that starts during embryonic cleavages and is still measurable postimplantation.

**Figure 4.**
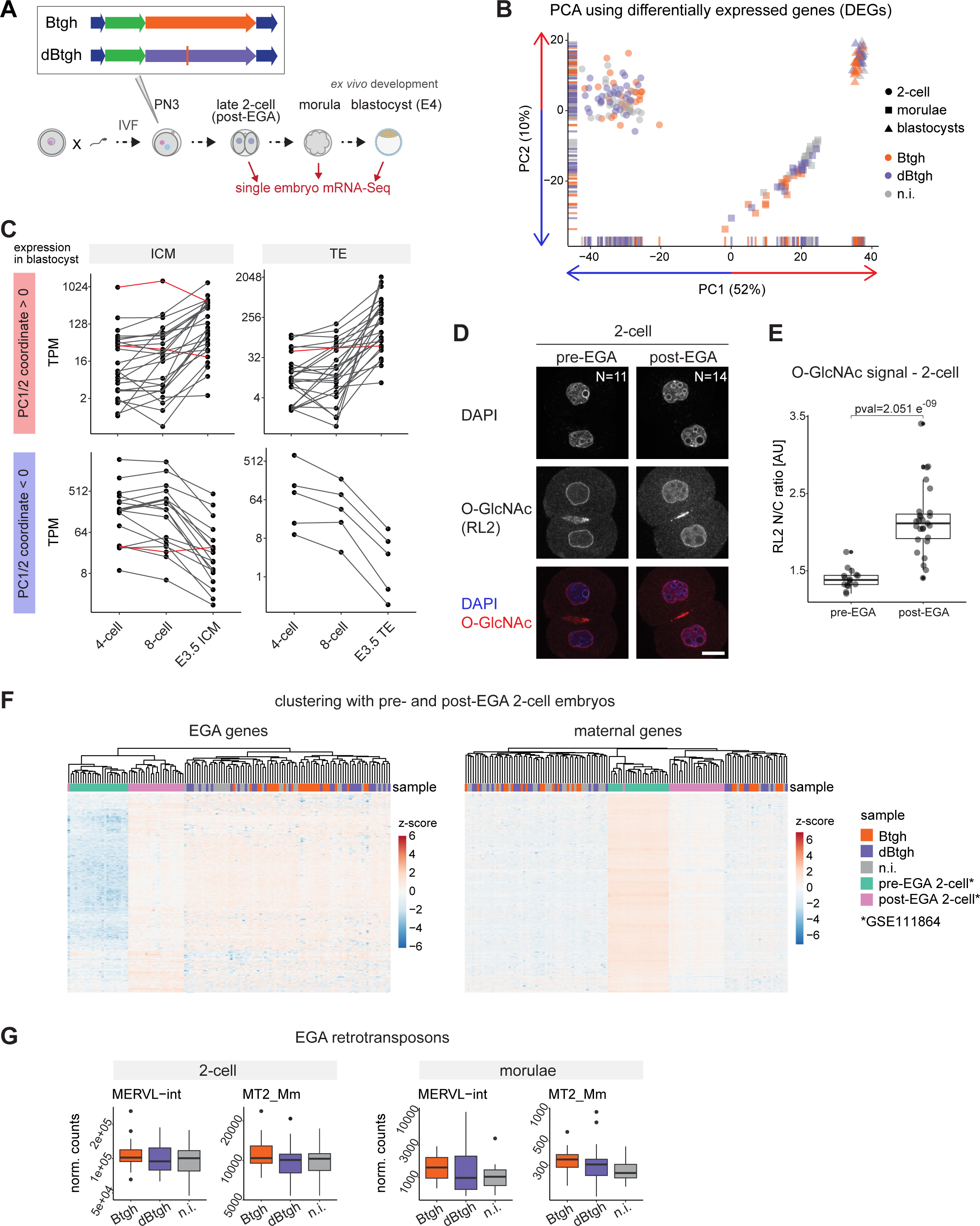
Nuclear O-GlcNAc is dispensable for EGA. (A) Experimental design for results in Figures 4 and 5. Zygotes were generated through IVF and injected 2 h later with the mRNA of Btgh or dBtgh or left uninjected. The three groups of embryos were cultured ex-vivo until the post-EGA 2-cell, morula and blastocyst stages and collected for single embryo mRNA-Seq. (B) PCA of single embryos’ transcriptomes at the three preimplantation stages, using all differentially expressed genes (DEGs) in O-GlcNAc-depleted embryos (adj. p-value < 0.05 in the Btgh vs. dBtgh or Btgh vs. non-injected comparison at any stage). The variance explained by each principal component is indicated in parentheses. N = 85 2-cell embryos, 56 morulae and 48 blastocysts. (C) Transcripts Per Million (TPM) across cleavage stages (mRNA-Seq data from GSE66582^32^ and^33^ GSE76505) of the 50 and 30 genes mostly contributing to, respectively, the variance explained by PC1 and PC2. Genes were divided based on their positive (upper row) or negative (bottom row) coordinates on the respective PC, and they were plotted in two groups based on the E3.5 blastocyst tissue (ICM: inner cell mass; TE: trophectoderm) where they showed higher TPM. (D) Immunofluorescence staining of O-GlcNAc (RL2 antibody) in pre- and post-EGA 2-cell embryos generated through IVF. One z-plane is shown for each embryo. Embryos were mounted on coverslips and imaged using a scanning confocal microscope. DNA was stained with DAPI. Scale bar indicates 20 μm. The total number of embryos imaged per stage is indicated. (E) Quantification of the nuclear to cytosolic (N/C) ratio of O-GlcNAc signal in single blastomeres of the 2-cell embryos shown in (D). Data are from two replicates of the IVF experiment, which were pooled in one single plot. P-value was computed using unpaired Wilcoxon rank sum exact test. (F) Heatmap of the three experimental groups of single 2-cell embryos from this study together with pre-EGA and post-EGA 2-cell embryos generated through ICSI (GSE111864)^45^. All samples were pooled in one DESeq2 dataset, the data normalized using DESeq2 and log2 transformed. Then, all strictly EGA genes (left) and all strictly maternal genes (right) with mean of DESeq2-normalized counts ≥ 10 in the combined 2-cell dataset (this study plus GSE111864) were used to build the heatmap. Heatmap values are scaled by rows and both genes and samples are clustered based on Pearson correlation. (G) DESeq2-normalized counts of two types of retrotransposons associated with mouse EGA. MERVL-int denotes the full-length MERVL element, MT2_Mm the MERVL 5’ LTR. Padj = adj. p-value computed using DESeq2 Wald test and corrected for multiple testing using the Benjamini and Hochberg method. Y-axes ticks are in log10 scale.

The nuclear O-GlcNAc signal in the preimplantation embryo showed a sharp increase specifically at the 2-cell stage (Figure 1B). Strikingly, this does not happen at the time of syngamy but during the 2-cell stage itself, coincidently with EGA (Figure 4D,E). This observation, together with the modification of RNA PolII CTD by O-GlcNAc^18^, prompted us to investigate whether nuclear O-GlcNAc depletion alters the process of EGA. To this end, we clustered the 2-cell transcriptomes from our dataset with published mRNA-Seq data of single 2-cell embryos collected 18 h (pre-EGA) and 28 h (post-EGA) after intracytoplasmic sperm injection (ICSI)^45^. We defined “strictly EGA’’ genes by intersecting published EGA genes^46^ and genes significantly upregulated (adj. p-value < 0.05 and log_2_FC > 1) from 18 h to 28 h in 2-cell embryos; we defined “strictly maternal” genes as the ones significantly downregulated (adj. p-value < 0.05 and log_2_FC < −1) from 18 h to 28 h in 2-cell embryos. Notably, both the activation of embryonic genes and the degradation of maternal transcripts clustered the samples from our experiments with post-EGA 2-cell embryos, with no additional clustering due to O-GlcNAc removal (Figures 4F and S3A). Therefore, the transcriptional maternal-to-zygotic transition is overall unaffected by nuclear O-GlcNAc depletion. In good agreement, the expression of known EGA markers (*Sp110*, *Zfp352*, *Zscan4c*, *B020004J07Rik*, *Arg2*, *Tcstv3*) was either unchanged or increased upon O-GlcNAc depletion (Figure S3B).

A spike in the expression of the MERVL class of LTR retrotransposons is a unique feature of the 2-cell stage transcriptome^47^. Both full-length MERVL (MERVL-int) and MT2 5’ LTRs (MT2_Mm) are associated with EGA^48^ and might be necessary for this process^49,50^. The two types of MERVL transcripts were equally expressed or slightly upregulated between Btgh-injected and control embryos (Figure 4G). Furthermore, we found no differentially expressed retrotransposons in O-GlcNAc-depleted 2-cell embryos (Figure S3C). O-GlcNAc-depleted morulae showed a few significantly deregulated retrotransposons (adj. p-value < 0.1; Figure S3C), but the difference in expression was very low in magnitude (0.1 < log_2_FC < 0.4). Most of the retrotransposons deregulated in morulae are dynamically expressed across preimplantation stages, thus their level in O-GlcNAc-depleted embryos could derive from the developmental delay. For example, the significantly upregulated MTA_Mm-int is an ERVL-MaLR element typical of the 1-cell embryo^48^ which should be downregulated between the 4-cell and 8-cell stage (Figure S3D). The trend of sustained upregulation of MERVL-int and MT2-Mm elements up to the morula stage after depletion of O-GlcNAc (Figure 4G) complies with this interpretation.

Taken together, our results demonstrate that EGA - including the upregulation of EGA-specific retrotransposons - and the degradation of maternal transcripts are globally unaffected by nuclear O-GlcNAc removal. Hence, we conclude that the developmental delay observed in the morula transcriptome starts after the completion of the maternal-to-embryonic transition.

### Misregulation of mitotic and translation-related genes in nuclear O-GlcNAc-depleted embryos

Although the depletion of nuclear O-GlcNAc did not measurably impact EGA, post-EGA 2-cell embryos displayed a widespread change in gene expression of low magnitude (Figure 5A). GSEA did not identify any specific upregulated pathway, while downregulated genes tend to be involved in the process of translation (Figure 5B). Accordingly, genes encoding ribosomal proteins (*Rpl12*, *Rpl11*, *Rps5*, *Rps15a*, *Rps26*, *Rps2*, *Rpl10*) as well as factors of translation initiation and mRNA metabolism (*Eif4b*, *Eif3a*, *Ythdf2*, *Larp4*) were significantly downregulated after O-GlcNAc depletion (Figures S4A and S4B). When averaging the expression of all ribosomal protein genes^51^, Btgh-injected embryos showed a mild but significant downregulation (p-value = 0.007; Figure 5C). The embryonic translational machinery is robustly transcriptionally activated during preimplantation development, particularly at the 2-cell stage^52^, as shown by the coordinated increase in the expression of ribosomal protein genes (Figure S4C). Thus, a subtle developmental delay already starting in post-EGA 2-cell embryos upon O-GlcNAc depletion could potentially explain the downregulation of translation-related genes. We tested this hypothesis by plotting the expression of the genes (not including those directly involved in translation) with a higher than 2-fold increase between the 2- and 4-cell stages. O-GlcNAc-depleted 2-cell embryos showed a lower average expression of these genes (p-value = 0.035; Figure 5D) albeit this did not correlate with the lower expression of ribosomal protein genes (Figure S4D). Therefore, we cannot establish whether the lower level of translation-associated gene counts is a cause or a consequence of a putative developmental delay already starting at the late 2-cell stage. Nevertheless, the earliest transcriptional phenotype that we measure after O-GlcNAc removal in the developing embryo is the dampening of the activation of the translational machinery.

**Figure 5.**
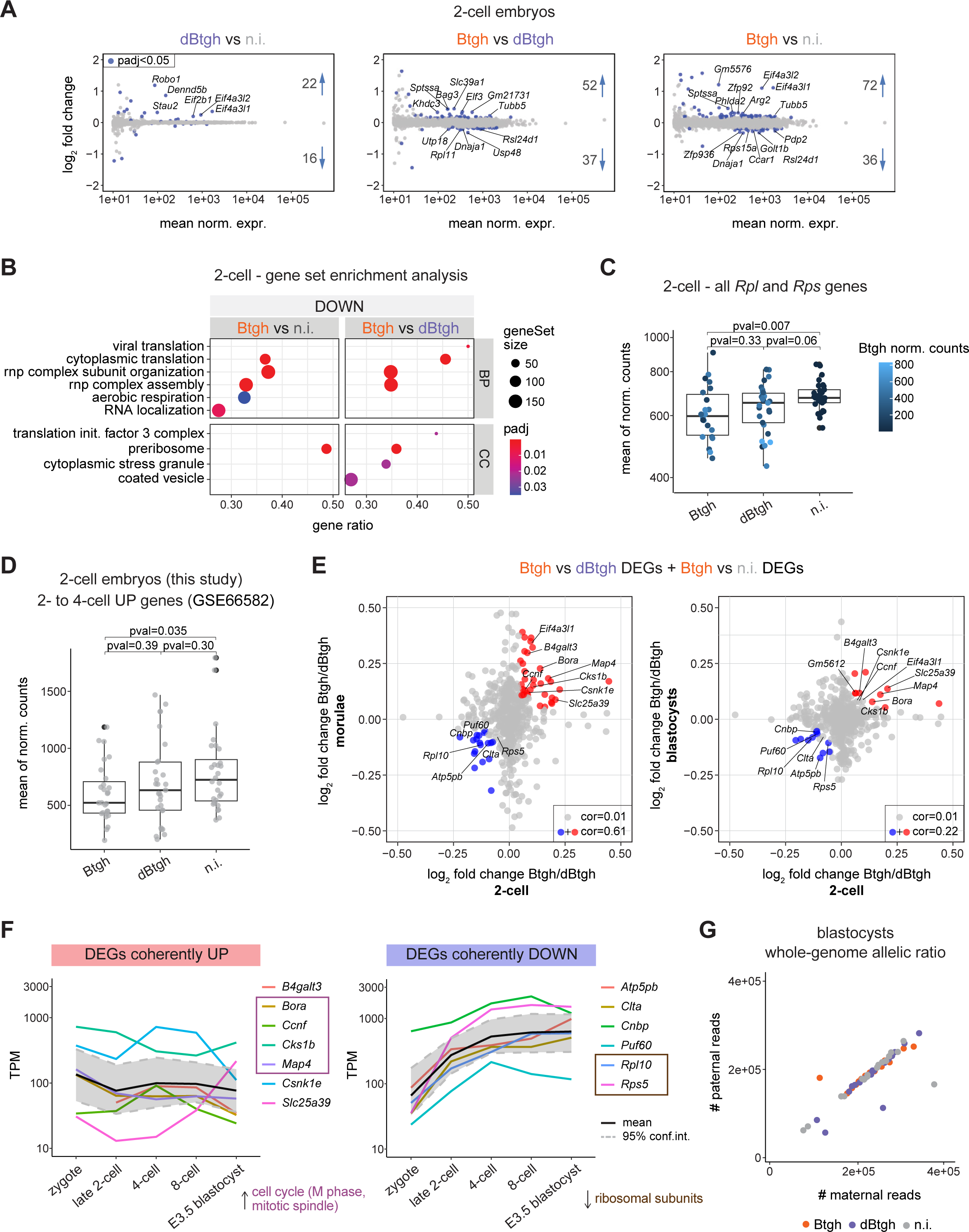
Misregulation of mitotic and translation-related genes in nuclear O-GlcNAc-depleted embryos. (A) MA-plots from DESeq2 differential gene expression between the indicated experimental groups of 2-cell embryos. Only genes with mean of DESeq2-normalized counts ≥ 10 are shown. All genes with adj. p-value < 0.05, any log2FC are colored, and their number is indicated. Genes standing out are labeled. (B) Gene set enrichment analysis of gene expression changes in nuclear O-GlcNAc-depleted 2-cell embryos versus the indicated controls. The 10 significant gene ontology (GO) terms with the highest Normalized Enrichment Score are shown for Biological Process (BP) and Cellular Component (CC) GOs, ordered by gene ratio. Only one upregulated term (*trophoblast giant cell differentiation*) was found among these 20 and it is not shown in the plot. The size of dots is proportional to the number of total genes of a GO term. Gene ratio = fraction of total genes of the GO term which are concordantly changing between the two conditions. (C) Single-embryo mean of DESeq2-normalized counts of all Mus musculus ribosomal protein genes from the Ribosomal Protein Gene Database^51^. Y-axis ticks are in log2 scale. Dots are colored based on DESe-q2-normalized counts of the Btgh/dBtgh RNAs. (D) Single-embryo mean of DESeq2-normalized counts of genes which are strongly upregulated between the post-EGA 2-cell and the 4-cell stage in publicly available mRNA-Seq data (GSE66582^32^, Methods). One outlier non-injected embryo is excluded from both plot and statistical test. (C,D) P-value was computed using Wilcoxon rank sum exact test. (E) Comparison of transcriptional deregulation across preimplantation stages, shown as a scatterplot of the log2 fold changes (between Btgh-injected and dBtgh-injected embryos) measured at each stage for all differentially expressed genes (DEGs) in O-GlcNAc-depleted embryos (adj. p-value < 0.05 in the Btgh vs. dBtgh or Btgh vs. non-injected comparison at any stage; DEGs with DESeq2-normalized counts < 10 at any stage were excluded from the plot). In each plot, DEGs with abs(log2FC) > 0.05 at both stages and abs(log2FC) ≥ 0.1 in at least one of the two stages are colored (coherently deregulated between two stages); among those, the ones with abs(log2FC) > 0.05 at all stages are labeled (coherently deregulated at all stages). cor = Pearson correlation. (F) Expression dynamics of DEGs coherently deregulated at all stages (labeled in (E); the two pseudogenes were excluded) throughout preimplantation development (mRNA-Seq data from GSE66582^32^ and GSE76505^33^). The two biological replicates per stage were averaged. The mean for all genes is drawn, as well as the 95% confidence interval, computed using basic nonparametric bootstrap (R function ‘mean.cl.-boot’). Y-axes shows Transcript Per Million (TPM) and ticks are in log10 scale. (G) Parental contribution to the whole autosomal genome in single Btgh/dBtgh-injected and non-injected FVB/PWD F1 blastocysts, measured as the total number of maternal (FVB) and paternal (PWD) mRNA-Seq reads. Outliers from the diagonal display allelic imbalance, thus represent potential aneuploid embryos.

At later preimplantation stages, the magnitude of gene expression changes due to O-GlcNAc depletion remained overall modest and the highest number of DEGs was found in the morulae (Figure S4E). When we compared the GSEA of morulae and blastocysts, we found several biological processes affected in both stages (Figure S4F). Downregulated genes participate in various essential cellular functions, including the mitochondrial respiratory chain complex, intracellular and membrane transport, the actomyosin cytoskeleton, the proteasome and DNA compaction. The downregulation of genes participating in aerobic respiration, as well as vesicle coat, already appeared in O-GlcNAc-depleted 2-cell stage embryos (Figure 5B). On the other hand, upregulated genes in morulae and blastocysts were mostly related to the metaphase-to-anaphase cell cycle transition (Figure S4F). We expect that a consistent fraction of the transcriptional differences observed at later preimplantation stages is a consequence of the developmental delay caused by O-GlcNAc depletion and described above (Figure 4B). For example, the direction of change shown by ribosome-related GOs in morulae and blastocysts from our dataset is the mirror opposite of their direction at corresponding stages in publicly available data (Figure S4G).

Next, we analyzed the dynamics of DEGs (adj. p-value < 0.05 in Btgh-injected embryos versus either of the two controls at any stage) across the three preimplantation stages. DEGs were poorly overlapping (Figure S4H) and their log_2_FC was uncorrelated between stages (Figure 5E), as expected from three developmentally distant time points with stage-specific transcriptomes.

However, a set of DEGs was persistently and coherently deregulated from the 2-cell embryo to the morula, and a smaller subset even up to the blastocyst (Figure 5E). Most of the coherently upregulated genes (7 in total) are involved in the transition to the mitotic phase of the cell cycle (*Ccnf*, *Bora*, *Cks1b*) or in the mitotic spindle (*Bora*, *Map4*), but they also include the mitochondrial glutathione transporter *Slc25a39*^53^, *B4galt3*, one of the 7 beta-1,4-galactosyltransferases responsible for the synthesis of complex-type N-linked oligosaccharides^54^, and Casein Kinase 1 Epsilon (*Csnk1e*), a central component of the circadian clock^55^. The constantly downregulated genes (6 in total) include two ribosomal proteins (*Rpl10* and *Rpl5*), Poly(U) Binding Splicing Factor 60 (*Puf60*), involved in pre-mRNA splicing^56^, a subunit of mitochondrial ATP synthase (*Atp5pb*), Clathrin Light Chain A (*Clta*), which coats intracellular vesicles, and CCHC-Type Zinc Finger Nucleic Acid Binding Protein (*Cnbp*), involved in transcriptional repression by binding the sterol regulatory element (SRE)^57^. Notably, when we looked at the RNA level of coherently deregulated genes across preimplantation stages in unperturbed embryos, we found that the downregulated ones are normally gradually upregulated throughout preimplantation development, up to 10-fold (Figure 5F, bottom panel). The highly similar magnitude of change across stages after Btgh injection (Figure 5E) suggests that their expression is continuously dampened in the absence of O-GlcNAc. It is also possible that their persistent transcriptional downregulation is contributed by the developmental delay caused by O-GlcNAc depletion, in the case of such a delay already starting in post-EGA 2-cell embryos, a possibility that we could not exclude (Figure 5D).

On the other hand, coherently upregulated genes showed more complex expression dynamics in unperturbed preimplantation embryos (Figure 5F, upper panel), thus their misregulation is unlikely a consequence of a delayed transcriptome but rather an acute response to the absence of O-GlcNAc. The upregulation of mitotic genes could indicate a dysfunction in chromosome segregation, leading to aneuploidy. To test this possibility, we quantified paternal versus maternal allelic contribution to the autosomal genome in single blastocysts depleted of nuclear O-GlcNAc. This was achieved using F1 hybrid embryos from evolutionary distant parental mouse strains (female FVB/NCrl x male PWD/Ph; Figure S4I). This analysis did not reveal a higher incidence of maternal/paternal allelic imbalance after nuclear O-GlcNAc depletion (Figure 5G).

In summary, the data revealed that nuclear O-GlcNAc depletion from the late zygote causes a developmental delay that starts after the completion of the maternal-to-embryonic transition. Around this time, the normal upregulation of the genes involved in translation is reduced when O-GlcNAc homeostasis is perturbed. In addition, spindle checkpoint genes are upregulated throughout preimplantation stages, although this is not associated with major consequences on mitosis.

## Discussion

O-GlcNAc, the only known form of reversible intracellular glycosylation in animals, is found on thousands of mammalian proteins. The embryonic lethality caused by maternal *Ogt* loss-of-function implies that O-GlcNAc or OGT are required for oocyte maturation, fertilization or post-fertilization before the expression of a paternal functional copy of the gene. To our knowledge, the function of the O-GlcNAc modification in the early embryo has not been investigated, nor its dynamics. In this study, we started by characterizing for the first time the localization and abundance of O-GlcNAc and its writer and eraser enzymes across preimplantation development. This new knowledge was fundamental to rationalize the strategy to functionally assess the role of O-GlcNAc in the early embryo.

First, we discovered that the localization pattern of OGT and O-GlcNAc is uncoupled in the first phases of development. In particular, in the zygote the O-GlcNAc transferase is strikingly enriched in the paternal pronucleus specifically. Because the payload of zygotic OGT proteins comes from the oocyte, the localization to the paternal pronucleus implies an active process, which could potentially have a functional significance; for instance, the need to exclude the protein from a certain compartment or to concentrate it on the paternal chromatin. Of the handful of factors for which such an asymmetrical pattern has been reported, a notable one is 5-hydroxymethylcytosine (5hmC). 5hmC is catalyzed by ten-eleven translocation (TET) enzymes, is enriched at the paternal pronucleus similarly to OGT and is believed to be associated with paternal DNA demethylation in the zygote^58–60^. Curiously, TETs are some of the most abundant OGT interactors^61^. The possibility of OGT function in the paternal pronucleus being related to zygotic DNA demethylation is intriguing, perhaps representing a missing player in this incompletely understood process^62^. Alternatively, OGT interaction with paternal chromatin (for example through TET enzymes) might exclude it from other cellular compartments as a way to tune O-GlcNAc concentration. It is important to bear in mind that embryonic cleavages are not accompanied by embryonic growth, hence O-GlcNAc homeostasis during this period might require lowering OGT effective concentration. This hypothesis fits well with our finding of a low abundance of full-length *Ogt* transcript at EGA, which indicates that, either through alternative splicing or other post-transcriptional mechanisms, the embryonic genome initially produces very little of the catalytically-competent *ncOgt* mRNA.

We showed that the subcellular pattern of O-GlcNAc is dynamically remodeled across developmental stages, with a remarkable increase in its nuclear abundance coincidently with EGA. This finding guided the design of our functional approach, consisting of the injection in the zygote of a nuclear-targeted bacterial homologue of OGA, Btgh. Our system based on enzymatic removal has three key features for addressing the function of O-GlcNAc PTM itself in early development: it does not interfere with OGT function; it cannot affect fertilization as observed upon OGA inhibition^26^; by targeting Btgh to the nucleus, we minimize the risk of inducing cell lethality or proliferation arrest, which until now have obfuscated the role of O-GlcNAc in embryonic gene expression. In fact, the importance of O-GlcNAc in fundamental cellular processes such as the cell cycle has been described^14^ and could explain, at least in part, the lethality observed in any proliferating cell type where *Ogt* was knocked-out^6,63^ while the deletion of *Ogt* in post-mitotic neurons did not affect cell viability^64^.

Through our novel functional approach, we unexpectedly found out that the reduction of nuclear O-GlcNAc to undetectable levels does not interfere with the activation of the embryonic genome, nor with the gene networks responsible for the first embryonic lineage specification. This result was unanticipated because O-GlcNAc is believed to contribute to pluripotency and developmental gene expression^15,65^. Our finding stresses the importance of assessing the functional relevance of any given PTM *in vivo* in its physiological context.

Instead, the data showed that nuclear O-GlcNAc removal slowed down embryonic development, starting from cleavage stages and persisting after implantation. Notably, at the transcriptional level this was associated with the dampening of the 2-cell-specific activation of a set of genes whose regulation is closely linked with growth demand, namely translation-related genes, including many ribosomal protein (RP) genes. Cells dedicate a large fraction of their transcription to ribosome biogenesis, hence the need to adapt this effort to nutrient and energy conditions. In both yeast and mammalian cells, rRNA and RP gene transcription is regulated by nutrient sensors such as target of rapamycin (TOR) signaling^66^. At the same time, a coordinated ribosomal gene production is necessary to ensure proper growth. Thus, although our experiments cannot establish the directionality of the interplay between transcriptional downregulation of translation and the observed growth retardation, the two phenomena are interconnected. We can speculate about possible nuclear O-GlcNAc targets acting on both growth and ribosomal genes, for example factors of the mTORC signaling cascade itself. In the cytosol, an activator of mTORC, named Raptor, has recently been reported to signal glucose status via its O-GlcNAcylation^67^.

Together with ribosome-related genes, a set of genes, all linked to the mitotic phase of the cell cycle, is upregulated at all stages after O-GlcNAc removal. Notably, our profiling revealed that OGT is enriched on the oocyte meiotic spindle (Figure 1B). Moreover, a crosstalk between O-GlcNAcylation and phosphorylation has been reported for several proteins associated with mitotic progression^68^, thus it is possible that perturbing the balance between these PTMs can lead to a delay in cell division, uncoupled from major mitotic defects, as suggested by the normal cleavage and absence of aneuploidy. In this regard, it is worth noticing that the breakage of the nuclear envelope at mitosis results in Btgh free diffusion to the cytosolic compartment during this phase. Alternatively, a cell cycle delay might be an indirect response to the slowdown of translation rate. Of note, differences in the rate of protein turnover as well as in the duration of cell cycle during embryonic cleavages have been both associated with the species-specific pace of embryonic development, via independent mechanisms^69–71^.

Finally, because of the abundance of O-GlcNAc in the nucleus and the function of many of its targets in transcription, the overall mild transcriptional response upon its removal was surprising. With transcriptomics as our main readout, it is possible that we are missing other types of molecular phenotypes related to chromatin changes or post-transcriptional processes, including nuclear transport, which might be impacted given the enrichment of O-GlcNAc on the NPC^27,28^.

This first functional exploration of O-GlcNAc biology *in vivo* in the mammalian embryo demanded a global and unbiased perturbation approach. It is important to bear in mind that, with such an approach, it is difficult to establish a direct link between O-GlcNAc and the observed transcriptional and developmental phenotypes. This is partly due to the low input of the embryonic material which limits biochemical investigations, but mostly, it is intrinsic to the pleiotropy of O-GlcNAc, which has the ability to orchestrate a cellular response by acting simultaneously on many targets. The global nuclear removal of O-GlcNAc uncovered embryonic growth control as its most phenotypically relevant function. Taken together with the associated transcriptional response affecting translation and the cell cycle, this O-GlcNAc function could have a long-reaching significance, since both these processes have been proposed as the molecular basis of species-specific developmental speed^69–71^.

With embryonic development representing an extremely dynamic metabolic system, and the donor of O-GlcNAc being responsive to nutrients, this study should pave the way for a further dissection of this novel intriguing connection between nuclear glycosylation and embryonic growth.

## Acknowledgements

We thank the EMBL Rome Gene Editing and Embryology Facility for assistance with embryo implantation, particularly Tianwu Qiao. We also thank the EMBL Rome Laboratory Animal Resource facility for animal welfare and husbandry, the EMBL Rome Microscopy facility for technical support and the EMBL GeneCore for library preparation and RNA-sequencing. We are grateful to Francesco Tabaro and Charles Girardot for their invaluable help with data management. We are grateful to Robyn L. Ball, Vivek Philip, Hongping Liang, and Elissa J. Chesler of the Jackson Laboratory for the comprehensive FVB/PWD SNPs annotation. Furthermore, we thank Takashi Ishiuchi and Jamie Hackett for critical reading of the manuscript. This research was funded by the European Molecular Biology Laboratory.

## Author contributions

S.F., A.B and M.B conceptualized the study. S.F. designed, conducted most *in vivo* experiments and performed all the bioinformatics analyses. U.C. performed most of the confocal microscopy experiments for OGT and O-GlcNAc profiling across preimplantation stages. N.L. supported the in vitro fertilization and microinjections experiments. A.S. conducted the micro-dissection of postimplantation embryos. S.F., M.B. and A.B. wrote the manuscript with input from all authors.

M.B. and A.B. jointly supervised the work.

## Declaration of interests

The authors declare no competing interests.

## STAR Methods

### STAR KEY RESOURCES TABLE

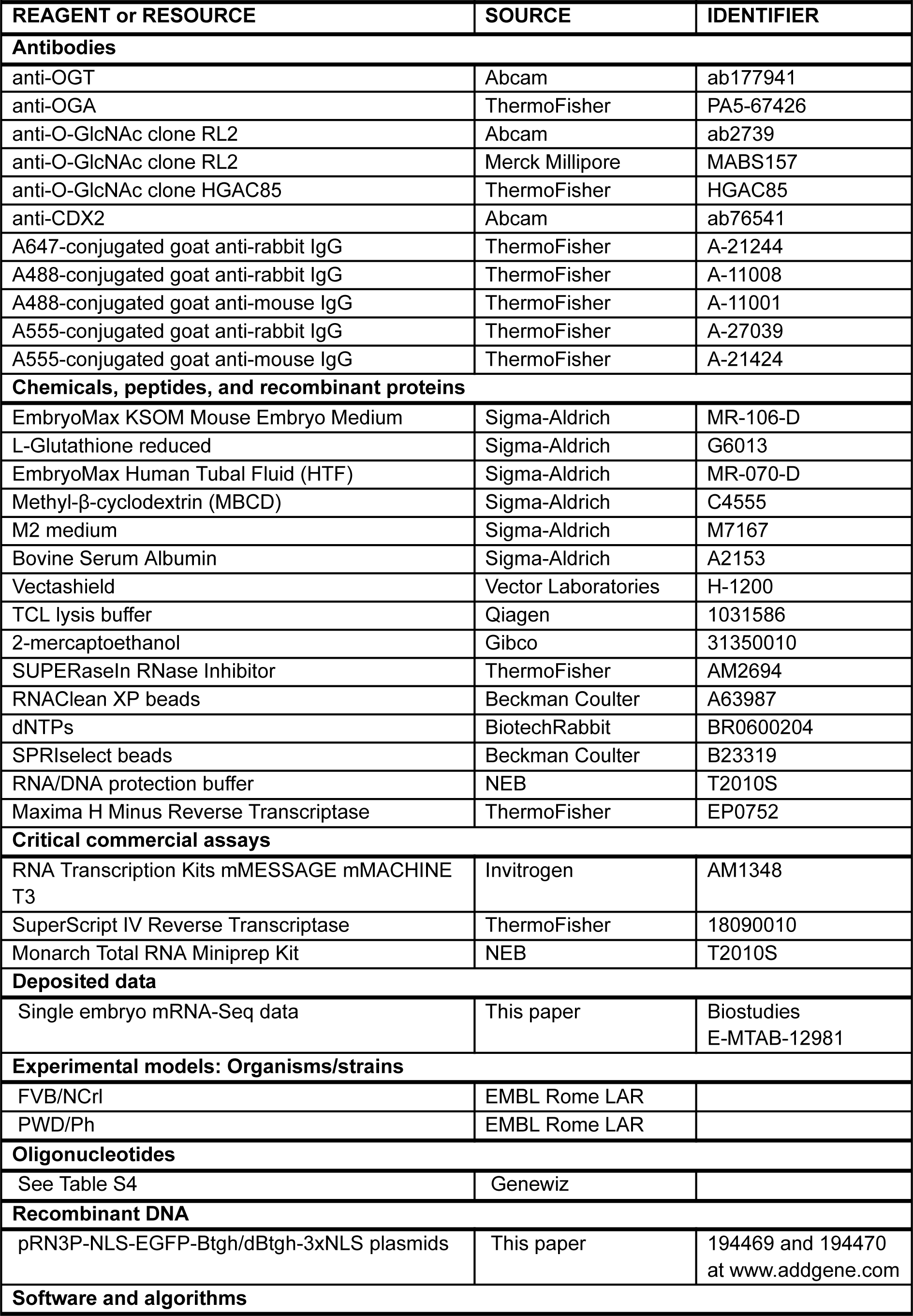
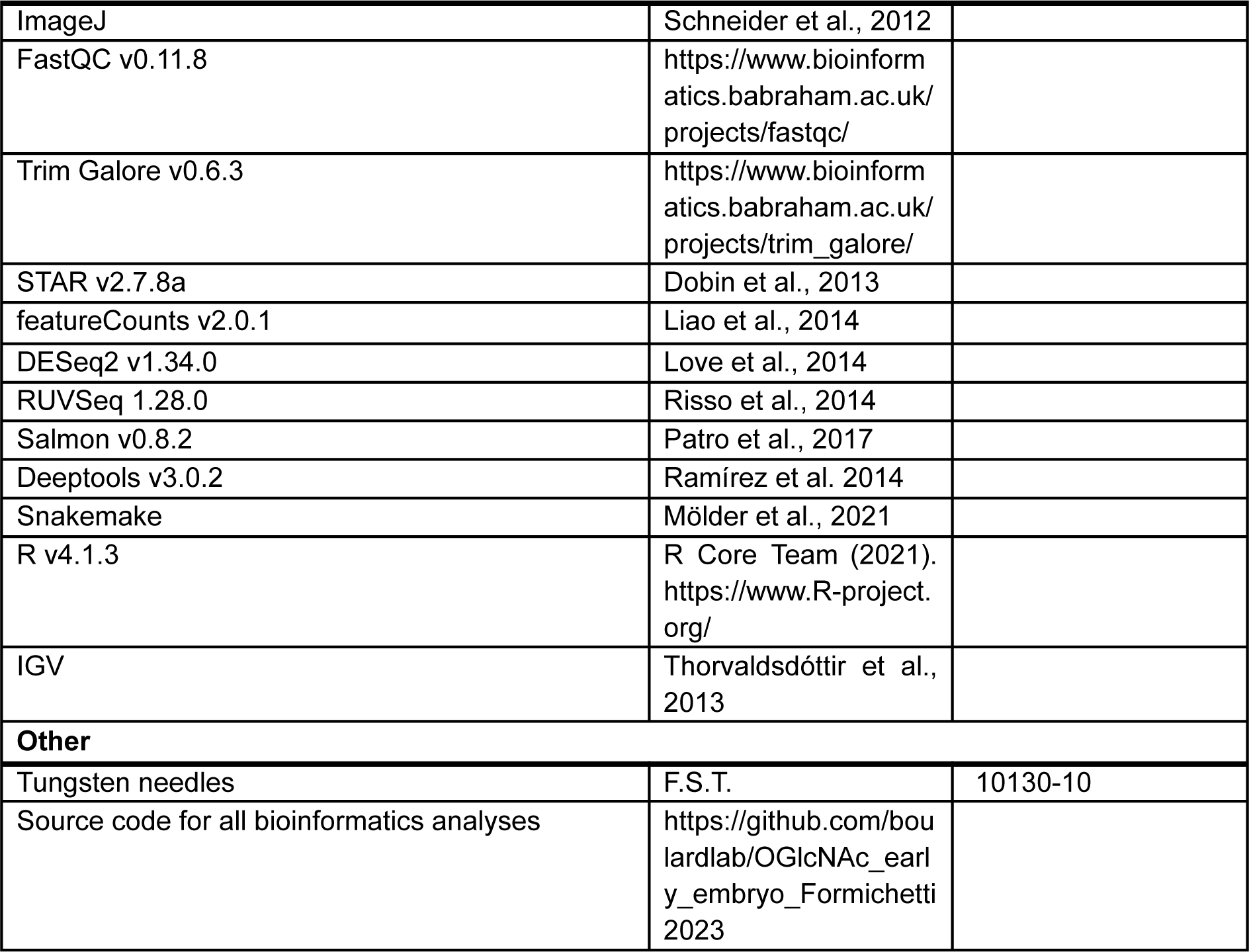

## RESOURCE AVAILABILITY

### Lead contact

Further information and requests for resources and reagents should be directed to and will be fulfilled by the lead contact, Matthieu Boulard.

### Materials availability

The plasmids generated in this study are available at www.addgene.org under accession numbers 194469 and 194470.

### Data and code availability

● The RNA-seq datasets generated in this study have been deposited in Biostudies and are available under the accession E-MTAB-12981.
● The previously published data used in this study are available under the accession numbers: GSE76505, GSE66582 and GSE111864.
● The source code of all bioinformatics analyses is available at: https://github.com/boulardlab/OGlcNAc_early_embryo_Formichetti2023

## EXPERIMENTAL MODEL AND SUBJECT DETAILS

### Animal care and strains

All procedures involving mice were handled in compliance with the rules and regulations of the Institutional Animal Care and Use Committee (IACUC) under protocol number 21-012_RM_MB, as well as with Italian Law (DL 26/2014, EU 63/2010) under protocol numbers 17/2019-PR to C.L. and 598/2023-PR to M.B. (Ministero della Sanità, Roma, Italy). Mice were housed in the pathogen-free Animal Care Facility at EMBL Rome on a 12-hours light-dark cycle in temperature and humidity-controlled conditions with *ad libitum* access to food and water. Female FVB/NCrl were used throughout the study; male FVB/NCrl were used for all imaging and NGS experiments except for the blastocyst and E7 Smart-Seq experiments, for which PWD/Ph males were used.

### Preimplantation embryo culture

Embryos were cultured in a standard mammalian cell incubator (37°C, 5% CO_2_) in EmbryoMax KSOM Mouse Embryo Medium (Sigma-Aldrich #MR-106-D).

## METHOD DETAILS

### Cloning of the pRN3P-NLS-EGFP-Btgh-3xNLS plasmid

The cDNA encoding O-GlcNAc hydrolase (Oga, *Btgh84*) from Bacteroides was amplified from the plasmid described in Boulard *et al.*^37^ and cloned in the pRN3P-H3.3R8A-GFP plasmid using restriction enzymes NotI and SmaI, fused to enhanced GFP *mgfp5* and flanked by SV40 NLSs. pRN3P-H3.3R8A-GFP is a generous gift from Maria Elena Torres-Padilla^72^. The catalytically inactive (“dead”) *Btgh84^D^*^242^*^A^* (characterized in Dennis et al.^36^) was cloned the same way. Sequences of primers and synthetic double-stranded DNA (IDT) used for plasmids cloning are in Table S2.

### In vitro transcription of NLS-EGFP-Btgh-3xNLS Btgh mRNA

The plasmid was linearized by digestion with KpnI, purified and then used as a template for mRNA synthesis with mMESSAGE mMACHINE T3 (Invitrogen #AM1348). In vitro transcribed RNA was purified with RNA Clean & Concentrator Kit (Zymo Research) and eluted in H_2_O. The mRNA product was run on an RNA ScreenTape with TapeStation 4150 (Agilent) to verify the presence of one main band of the correct size, the concentration was measured with NanoDrop 2000 spectrophotometer (ThermoFisher) and working concentration aliquots were made by diluting the mRNA in H_2_O. Aliquots were stored at −70°C until microinjection.

### In vitro fertilization (IVF)

In vitro fertilization was performed based on the published protocol, with minor modifications ^73^. Briefly, superovulation of 6-8 weeks FVB females was induced by hormonal stimulation (5 IU of PMSG and 5 IU of hCG 64 h and 16 h before collection, respectively), and cumulus-oocyte complexes were collected in KSOM media containing GSH (final concentration 10 mM; Sigma-Aldrich #G6013). Concomitantly, cauda epididymis and vasa deferentia from FVB or PWD males were dissected and sperm was gently squeezed out into capacitation media (HTF supplemented with MBCS at final concentration of 0.75 mM) (HTF: Sigma-Aldrich #MR-070-D; MBCD: Sigma-Aldrich #C4555), and allowed to swim up for 1 hour. Sperm was subsequently counted and cumulus-oocyte complexes were inseminated with 0.2M sperm in a 200 µL fertilization drop. Four hours after sperm addition, zygotes were cleaned from the surrounding cumulus cells and sperm by 5-6 washes in KSOM, then cultured in KSOM for 2 hours prior to microinjections or for 0 to 4 hours prior to fixation for immunostaining.

### Zygote microinjection

Two hours after the end of IVF, zygotes were transferred to an injection plate with M2 medium at 37 °C, and subjected to microinjections. Zygotes were microinjected with either pRN3P-NLS-EGFP-Btgh-3xNLS or pRN3P-NLS-EGFP-deadBtgh-3xNLS mRNA, both at 100 ng/µL (for the 2-cell RNA-Seq and IF experiment) or 300 ng/µL (for all other experiments). RNA injections were carried out using a Femtojet microinjector (Eppendorf) at 150 hPa pressure for 0.1 seconds, with 10 hPa compensation pressure (estimated microinjection volume of 10 pL). After the microinjections, zygotes were washed 3-4 times in KSOM and placed back into culture. In the case of Btgh/dBtgh injections, GFP fluorescence was verified the next day and GFP-positive (injected) 2-cell embryos were sorted from the non-injected ones and used for downstream imaging, developmental or transcriptomics analyses.

### Immunofluorescence staining of preimplantation embryos

MII oocytes (20h post-hCG), zygotes (0, 2 and 4 hours post-IVF, where 0 h is the end of the IVF), 2-cell embryos (pre-EGA: 19 h post-IVF, post-EGA: 28 h post-IVF), morulae (46 h post-IVF) and blastocysts (68 h post-IVF) were treated the same way, as in Bošković *et al.*^74^ with some modifications as follows. After two washes in M2 medium (Sigma-Aldrich #M7167), the zona pellucida was removed by a brief incubation in drops of warm Acidic Tyrode’s solution (Sigma-Aldrich), followed by other two M2 washes to neutralize the acid. The embryos were then washed once in PBS + 0.5% BSA, fixed in 4% PFA in PBS for 20 min at 37°C, permeabilized in 0.5% Triton X-100 for 20 min at 37°C and washed three times in PBS-T (0.15% Tween-20 in PBS). The epitope was then unmasked in 50 mM NH_4_Cl solution in H_2_O for 10 min at room temperature, followed by two additional PBS-T washes and then blocking - for 3 h at room temperature or overnight at 4°C - in BSA (Bovine Serum Albumin, Sigma-Aldrich #A2153) 3% in PBS-T. Primary antibody incubation was performed overnight at 4°C in the blocking solution, followed by three washes in PBS-T, re-blocking for 30 min at room temperature, three additional PBS-T washes, secondary antibody incubation for 1 to 2 h at room temperature in the blocking solution and three final PBS-T washes. The embryos were immediately mounted on coverslips in Vectashield (Vector Laboratories #H-1200) containing 4’-6-Diamidino-2-phenylindole (DAPI) or - where specified in the figure legend - in drops of 75% Vectashield in PBS, to preserve the 3D structure.

The primary antibodies used were: anti-OGT (abcam #ab177941), anti-OGA (ThermoFisher #PA5-67426), anti-O-GlcNAc clone RL2 (abcam #ab2739 and Merck Millipore #MABS157), anti-O-GlcNAc clone HGAC85 (ThermoFisher #HGAC85), anti-CDX2 (abcam #ab76541). Dilution of all primary antibodies was 1:200. Secondary antibodies used were: A647-conjugated goat anti-rabbit IgG (#A-21244), A488-conjugated goat anti-rabbit IgG (#A-11008), A488-conjugated goat anti-mouse IgG (#A-11001), A555-conjugated goat anti-rabbit IgG (#A-27039), A555-conjugated goat anti-mouse IgG (#A-21424) (all from ThermoFisher). Dilution of all secondary antibodies was 1:500. Fixed immunostained samples were imaged using a Nikon AX scanning confocal (using galvanometric mirrors) or - when specified in the figure legend - an X-light V3 Spinning disk confocal.

### Preimplantation embryo collection and library preparation for single embryo Smart-Seq

Single post-EGA 2-cell embryos (28 h post-IVF), morulae (48 h post-IVF) and blastocysts (68 h post-IVF) were collected in 5uL of 1x TCL lysis buffer (Qiagen #1031586) containing 1% (v/v) 2-mercaptoethanol (Gibco #31350010) and 0.5 U/µL of SUPERaseIn RNase Inhibitor (ThermoFisher #AM2694). RNA was separated from the surrounding material using RNAClean XP beads (Beckman Coulter #A63987) according to the manufacturer’s protocol for a 96-well plate and small volume reactions. After the last ethanol wash, the RNA was eluted from RNA beads in 8 µL of H_2_O, 3 µL of which were transferred to a new plate containing 1 µL of dNTPs (BiotechRabbit #BR0600204, 10 mM each) and 1 µL of 10 µM oligo-dT primer (5′–AAGCAGTGGTATCAACGCAGAGTACT30VN-3′). The plate was then sent to EMBL Gene Core Facility, which used a modified Smart-Seq2 protocol^75^ using SuperScript IV RT and tagmentation procedure previously described^76^ to prepare single-embryo full-length cDNA sequencing libraries. The retrotranscription reaction mix was as follows: 2 μL SSRT IV 5x buffer, 0.5 μL 100 mM DTT, 2 μL 5 M betaine, 0.1 μL 1 M MgCl2, 0.25 μL 40 U/μL RNAse inhibitor, 0.25 μL SSRT IV, 0.1 μL 100 uM TSO, 1.15 μL RNase-free H_2_O; with thermal conditions: 52°C 15 min, 80°C 10 min. cDNA was generated using 18 PCR cycles. The cDNA cleanup (0.6x SPRI ratio; SPRIselect beads: Beckman Coulter #B23319) was carried out omitting the ethanol wash steps and the elution volume was 13 μL of H_2_O. For tagmentation, the sample input was normalized to 0.2 ng/uL. After tagmentation and PCR, 2 μL of each sample were pooled in a single tube before the final clean up using 0.7x SPRI ratio. The pool was sequenced in one run (40 bp paired-end mode) on the Illumina NextSeq 500 sequencer. See Table S3 for the number of pooled embryos and average number of reads obtained per embryo at each stage.

### Surgical transfer of embryos for *in utero* development, E7 dissection and preparation for Smart-Seq

CD1 females 7-8 weeks of age in estrus were mated to vasectomized males, and the day of the vaginal plug was considered to be 0.5 day of pseudopregnancy. Microinjected zygotes were kept in culture overnight. The morning after (the day of vaginal plug), 15-20 embryos from each experimental group were surgically transferred into the oviduct of each pseudopregnant female. Six days later (seven days after IVF, for which reason we consider the embryos to be embryonic day 7), the foster mother was sacrificed and the embryos removed from their implantation sites and dissected in PBS with 10% FBS (PAN/Biotech). Embryos were cleaned by removing Reichardt’s membrane and surrounding maternal cells using 0.01 mm Tungsten needles (F.S.T. #10130-10), then imaged with trans-illumination on Leica M205C dissection microscope with fixed objectives (10×16) and photographed. For Smart-Seq, embryos were bisected with a glass needle along the epiblast, extraembryonic ectoderm (ExE) boundary, then the two halves collected in 200 µL of 1x RNA/DNA protection buffer (NEB #T2010S) in PBS, snap frozen and stored at −70°C until RNA extraction. RNA from single embryonic halves was extracted using Monarch Total RNA Miniprep Kit (NEB #T2010S), following manufacturer’s instructions for mammalian whole blood with some modifications: 5 µL of Proteinase K were added to each thawed sample, followed by incubation at room temperature for 15 min. RNA was eluted with 25 µL of 1x TE buffer. For a balanced sequencing design, sex genotyping was performed using cDNA made from 2 µL of each sample. cDNA was synthesized with 100 U of Maxima H Minus Reverse Transcriptase (ThermoFisher #EP0752), using random hexamers and following manufacturer’s instructions. Sex determination was achieved by cDNA PCR genotyping using three pairs of primers, against the mRNA of *Xist*, *Ddx3y* and *Eif2s3y* genes, respectively (see Table S4 for primer sequences).

The quality of the RNA was checked on an RNA High Sensitivity ScreenTape with TapeStation 4150 (Agilent) and only good quality samples were used by for the Smart-Seq2 library preparation described above and sequencing on a NextSeq 500 (40 bp paired-end mode).

### Analysis of microscopy images

All steps were performed using ImageJ^77^ unless specified. For the quantification of O-GlcNAc signal in 2-cell embryos, the z-plane with the largest nuclear area and highest intensity of DAPI signal was selected for each blastomere nucleus. Then, for Figure 2C: i. a measuring line was drawn across each nucleus and the RL2 signal intensity recorded; ii. all single-nucleus measures were used to generate a line plot in R using function ‘geom_smooth’ with ‘gam’ method. For Figure 4E: i. an ellipse was drawn inside the cytosol and nucleus of each blastomere and the mean RL2 signal intensity recorded for both compartments; ii. a custom script in R was used to compute the N/C intensity ratios.

The size of E7 embryos was measured from microscopy images as follows. A line was drawn following the contour of each embryo (including epiblast and trophectoderm, excluding the ectoplacental cone; as shown in Figure S2C, upper part), and the area delimited by the line was recorded. One litter from dBtgh-injected embryos which was a clear outlier (all Mid-Streak or later stages) was excluded from the resulting plot (Figure 3C).

### Single embryo Smart-Seq data analysis for single copy genes

The analysis pipeline was performed using Galaxy^78^ to obtain the table of gene counts. Briefly, the quality of reads was analyzed using FastQC v0.11.8 (https://www.bioinformatics.babraham.ac.uk/projects/fastqc/). Reads were trimmed from adapters and low-quality 3’-end nucleotides using Trim Galore v0.6.3 (https://www.bioinformatics.babraham.ac.uk/projects/trim_galore/) with default parameters for paired-end libraries (-q 20--stringency 1-e 0.1--length 20 –paired), before mapping them using STAR v2.7.8a^79^ and default parameters for paired-end reads to the GRCm38 mouse genome containing the NLS-GFP-Btgh-3xNLS transgene. Gene counts were obtained from the bam files using featureCounts (subreads v2.0.1)^80^, with default parameters for paired-end reads and counting fragments instead of reads. The gene counts were used in a custom Rmd script as input for DESeq2 v1.34.0^81^. The test used for statistical significance was the Wald test and the significance cut-off for optimizing the independent filtering was 0.05.

For the 2-cell embryo and blastocyst datasets, which were the result of more than one embryo generation and collection, batch effect had to be considered when testing for differential expression using DESeq function. To this aim, package RUVSeq^82^ was used to compute the factors of unwanted variation, with function ‘RUVs’ and k=3. The 3 factors of unwanted variation computed with RUVSeq were used in ‘DESeq’ formula *∼ W1 + W2 + W3 + condition*.

Whenever MA-plots are shown, the log_2_ fold changes are shrunken using the ‘ashr’ method^83^. Before PCA and differential expression analysis, a few filtering steps were applied to all datasets: embryos with < 10^6^ reads and outlier embryos in a scatterplot of mitochondrial DNA (mtDNA) gene expression versus percentage of reads mapping to ribosomal DNA (rDNA) were removed as low-quality samples. In the blastocyst dataset, the outliers in Figure 5G (aneuploid embryos) were additionally removed; for E7, those halves showing contamination from the complementary tissue based on the expression of markers for epiblast (*Map7d3*, *Pdzd4*, *Uchl1*) and extraembryonic ectoderm (*Gjb3*, *Gm9*, *Wnt6*) were excluded from all analyses. The exact number of embryos filtered at each step is in Table S3.

### Analysis of publicly available mRNA-Seq data

For the analysis of datasets GSE66582 (Wu et al., 2016) and GSE76505 (Zhang et al., 2017), both transcript counts and bigwig files were obtained using Galaxy^78^. In detail, the quality of the reads was analyzed using FastQC, then reads trimmed from adapters and low-quality 3’-end nucleotides using Trim Galore v0.6.3, first with default parameters and automatic adapter detection, then by specifically removing Clontech SMART CDS Primer II A 25 nt sequence (5’-AAGCAGTGGTATCAACGCAGAGTAC-3’) from both forward and reverse reads. For gene expression analysis, reads were mapped to Gencode vM25 (GRCm38.p6) transcript sequences using Salmon v0.8.2^84^ with default parameters for unstranded paired-end libraries. The output transcript counts were used in a custom Rmd script as input for gene-level summarization using tximport^85^. Gene-level TPM values were extracted from tximport ‘abundance’ matrix. To find genes upregulated between late 2-cell and 4-cell embryos (Figure 5D) we used DESeq2; the test used for statistical significance was the Wald test, the significance cut-off for optimizing the independent filtering was 0.05 and the log_2_ fold changes were shrunken using the ‘ashr’ method^83^. Then, genes were selected if: i. 2-cell TPM > 1; ii. 4-cell TPM > 3rd quartile of expression among genes with TPM > 1; iii. 4-cell/2-cell log2FC > 2. The only *Rpl* gene selected by these criteria was excluded.

For the inspection of reads coverage and the analysis of alternative splicing, fastq files from more runs of the same sample were concatenated before trimming, then mapping was performed using STAR v2.7.8a^79^ in single-sample 2-pass mode for higher accuracy, after genome indexing optimized to read length (--sjdbOverhang set to 100). Bigwig files were obtained with deeptools v3.0.2, normalizing to bins per million. The sashimi plots (Figure S1D) were generated using ggsashimi^86^ with density overlay between biological replicates but aggregation of junction counts (parameters--aggr median_j--min-reads-coverage 3).

### Principal component analysis (PCA) of single embryo transcriptomes

All included PCAs were performed with function ‘prcomp’ on log_2_-transformed and DESeq2-normalized raw counts and only genes with mean DESeq2-normalized expression > 10 across the ‘prcomp’ input samples were used. For Figure S3A, among the EGA+maternal genes (defined in the Results section), the 200 genes with highest variance in GSE111864 were used to perform the PCA. For Figure 4B, all DEGs (adj. p-value < 0.05, any log_2_FC) from the Btgh versus dBtgh and Btgh versus non-injected comparison at any stage were used.

### Gene set enrichment analysis (GSEA)

Performed in a custom Rmd script using function ‘gseGO’ of R package ‘clusterProfiler’^87^ on all genes of the dataset with DESeq2-normalized mean across samples > 10, ranked by -log10(adj. p-value)*sign(log_2_FC). P-value cutoff was 0.05 and parameters set to default. Results were simplified based on adj. p-value after computing semantic similarity using ‘mgoSim’ function of R package ‘GoSemSim’^88^. Similarity cutoff was 0.6 for preimplantation datasets, 0.8 for the E7 epiblasts. For each figure, the first significant terms based on Normalized Enrichment Score (NES) are shown, ordered by GeneRatio.

### Analysis of transposable elements’ expression

The analysis was performed using a custom Snakemake v5.9.1 pipeline^89^, available at the GitHub repository linked below. In summary, quality of the fastq files was checked with FastQC v0.11.8 and reads were trimmed using Trim Galore v0.6.4 with default parameters (-q 20--stringency 1-e 0.1--length 20 –paired). Trimmed reads were aligned to GRCm38 mouse genome with STAR v2.7.5c^79^, with parameters recommended by Teissandier et al.^90^ for the analysis of transcripts derived from autonomous retrotransposons in the mouse genome:--outFilterMultimapNmax 5000--outSAMmultNmax 1--outFilterMismatchNmax 3--outMultimapperOrder Random--winAnchorMultimapNmax 5000--alignEndsType EndToEnd--alignIntronMax 1--alignMatesGapMax 350--seedSearchStartLmax 30--alignTranscriptsPerReadNmax 30000--alignWindowsPerReadNmax 30000--alignTranscriptsPerWindowNmax 300 –seedPerReadNmax 3000--seedPerWindowNmax 300--seedNoneLociPerWindow 1000. After alignment, using a custom script, we only kept reads from pairs where both mates were: i. completely included into a repetitive element; ii. not overlapping with gene bodies of Gencode vM25. Repeat Library 20140131 (mm10, Dec 2011) was used as repetitive elements annotation, after excluding “Simple repeats” and “Low complexity repeats”.

With the STAR parameters above, only one random alignment (the one with the highest alignment score) is reported for multimappers, preventing the precise quantification of single repetitive elements. Therefore, for each repName of the repetitive element annotation, often present in multiple copies in the genome, read counts were summarized using featureCounts (subreads v2.0.1)^80^, with parameters-p-B s 0--fracOverlap 1-M.

The downstream analysis of transposable elements’ expression was performed in a custom Rmd script. First, an analysis at family level (repFamily field in the Repeat Library annotation) was performed in order to identify samples with an important contamination from genomic DNA, since the latter can affect the quantification of retrotransposon transcripts. For this analysis, FPKM were computed for each repFamily and samples with high FPKM values for DNA transposon families were removed. Then, differential Expression of RNA transposons was tested at the RepName level, using DESeq2 v1.34.0^81^ and adding the value of the total sum of DNA FPKM as a confounding factor to DESeq formula (*∼ DNA_FPKM + condition*). The test used for statistical significance was the Wald test and the significance cutoff for optimizing the independent filtering was 0.05. Whenever MA-plots are shown, the log_2_ fold changes are shrunken using the ‘ashr’ method^83^.

### Allele-specific Smart-Seq data analysis

The analysis was performed using a custom Snakemake v5.9.1 pipeline^89^, available at the GitHub repository linked below. Firstly, the position and identity of all annotated Single Nucleotide Polymorphisms (SNPs) between the FVB/NCrl and PWD/Ph mouse strains was collected from GenomeMUSter, a resource of the Mouse Phenome Project (MPD, RRID:SCR_003212)^91^. The PWD genome is not included in the Mouse Genome Project, making the accuracy of SNPs annotation highly variable compared to the better annotated FVB strain. Therefore, only SNPs with confidence level = 1 for the FVB DNA base and ≥ 0.8 for the PWD DNA base were kept for all analyses and used to create a masked GRCm38 mouse genome with ‘maskfasta’ from the BEDTools suite^92^. After quality check with FastQC v0.11.8 and trimming using Trim Galore v0.6.4 with default parameters (-q 20--stringency 1-e 0.1--length 20 –paired), reads were mapped to the masked genome using STAR v2.7.5c^79^. Alignments parameters were the default ones, except for--alignEndsType EndToEnd and--outSAMattributes NH HI NM MD, both required by SNPsplit (see below). After keeping only uniquely aligned reads and removing duplicates with Picard Tools (http://broadinstitute.github.io/picard/) - as suggested in Castel et al.^93^ - reads were assigned to either the FVB or the PWD genome using SNPsplit (https://www.bioinformatics.babraham.ac.uk/projects/SNPsplit/). FVB reads and PWD reads were used for two separate instances of featureCounts (subreads v2.0.1)^80^ to obtain gene counts, with parameters-s 0-t ‘exon’-g ‘gene_id’-Q 0--minOverlap 1--fracOverlap 0--fracOverlapFeature 0-p-C. featureCounts output was then used in a custom Rmd for the analysis of ploidy, for which all gene counts for the FVB and PWD genome were summed for each embryo (excluding genes on sex chromosomes).

### Quantification and statistical analysis

All statistical tests used are specified in the figure legend of the corresponding result. The exact p-values are indicated in the figures, while adjusted p-values are indicated in figures only when <0.05 for single copy genes or <0.1 for retrotransposons and higher values are considered not significant. For boxplots, hinges correspond to first and third quartiles; median is shown inside; whiskers extend to the largest and smallest values no further than 1.5 * IQR from the hinge (IQR = inter-quartile range, or distance between the first and third quartiles); data beyond the end of the whiskers are plotted individually.

## Notes

### Competing Interest Statement

The authors have declared no competing interest.

