## Supplemental Tables for "Perturbing nuclear glycosylation in the mouse preimplantation embryo slows down embryonic growth"

### Table S1. Staging of E7 embryos based on widefield microscopy images

| | PS | PS/ES | ES | advanced ES | ES or MS | MS | Not determined | Tot determined | Pre-ES % | ES % | Adv./Post-ES % |
|---|---|---|---|---|---|---|---|---|---|---|---|
| **Btgh** | 2 | 1 | 8 | 0 | 2 | 2 | 9 | 15 | 13 | 60 | 27 |
| **dBtgh** | 2 | 4 | 7 | 2 | 1 | 4 | 11 | 20 | 10 | 55 | 35 |

Two-way Anova p-value = 0.2

PS = Primitive Streak, ES = Early Streak, PS/ES = between PS and ES, advanced ES = advance Early Streak, MS = Mid Streak, Not determined = staging unsure.

### Table S2. Primers and gBlock used for cloning the pRN3P-NLS-EGFP-Btgh/dBtgh-3xNLS plasmids

| Sequence type | Target | Primer direction | Sequence (5'-3') |
|---|---|---|---|
| Primer | Btgh-containing plasmid (Boulard et al., 2020) | forward | GGAAGCATCAGGCGGGCTGCAGAATAGTAAAGGAGAAGAACTTTTCACTGGAGTTGTCCCA |
| | | reverse | GAGAAACATTCTGTCCGCCTTTGTATAGTTCATCCATGCCATGTGTAATCC |
| Primer | NLS-mgfp5 gBlock | forward | CAAAGGCGGACAGAATGTTTCTCTCCAACCTCCAC |
| | | reverse | AGTGGTAACCAGATCCGCTCACACTTTCCGTTTCTTCTTAGGATCG |
| NLS-mgfp5 gBlock | | | GATCTGAATTCCTGCAGCCCGCCACCATGGCCTCTCCTAAGAAAAAGAGGAAAGTGGAAGCATCAGGCGGGCTGCAGAATAGTAAAGGAGAAGAACTTTTCACTGGAGTTGTCCCAATTCTTGTTGAATTAGATGGTGATGTTAATGGGCACAAATTTTCTGTCAGTGGAGAGGGTGAAGGTGATGCAACATACGGAAAACTTACCCTTAAATTTATTTGCACTACTGGAAAACTACCTGTTCCATGGCCAACACTTGTCACTACTTTC |

| | | | |
|---|---|---|---|
| | | | ACTTATGGTGTTCAATGCTTTTCAAGATACCCAGATCATATGAAGCGGCACGACTTCTTCAAGAGCGCCATGCCTGAGGGATACGTGCAGGAGAGGACCATCTTCTTCAATGACGACGGGAACTACAAGACACGTGCTGAAGTCAAGTTTGAGGGAGACACCCTCGTCAACAGGATCGAGCTTAAGGGAATCGATTTCAAGGAGGACGGAAACATCCTCGGCCACAAGTTGGAATACAACTACAACTCCCACAACGTATACATCATGGCCGACAAGCAAAAGAACGGCATCAAAGCCAACTTCAAGACCCGCCACAACATCGAAGACGGCGGCGTGCAACTCGCTGATCATTATCAACAAAATACTCCAATTGGCGATGGCCCTGTCCTTTTACCAGACAACCATTACCTGTCCACACAATCTGCCCTTTCGAAAGATCCCAACGAAAAGAGAGACCACATGGTCCTTCTTGAGTTTGTAACAGCTGCTGGGATTACACATGGCATGGATGAACTATACAAAGGCGGACAGAATGTTTCT |

Table S3. Details on the generation and filtering steps of the single embryo Smart-Seq datasets

| **Stage** | **# pooled embryos in sequencing run** | **Average # of reads per embryo** | **# pooled embryos in sequencing run, per group** | | **# embryos discarded for < $10^6$ reads** | **# embryos discarded for high rDNA/ mt gene counts** | **# embryos discarded for low Btgh expression** | **# embryos discarded for opposite tissue contamination** | **# embryos discarded for aneuploidy** | **# embryos for Diff. Expr. analysis** |
|---|---|---|---|---|---|---|---|---|---|---|
| 2-cell | 96 | $5.3*10^6$ | Btgh | 32 | 3 | 1 | 1 | Not applicable | Not assessed | 27 |
| | | | dBtgh | 32 | 2 | 0 | 1 | Not applicable | Not assessed | 29 |
| | | | n.i. | 32 | 3 | 0 | 0 | Not applicable | Not assessed | 29 |
| morula | 62 | $6.3*10^6$ | Btgh | 22 | 1 | 0 | 3 | Not applicable | Not assessed | 18 |

| | | | | | | | | | | |
|---|---|---|---|---|---|---|---|---|---|---|
| | | | dBtgh | | 18 | 1 | 0 | 1 | Not applicable | Not assessed | 16 |
| | | | n.i. | | 22 | 0 | 0 | 0 | Not applicable | Not assessed | 22 |
| blastocyst | 62 | $6.1* 10^6$ | Btgh | | 19 | 0 | 2 | 0 | Not applicable | 3 | 14 |
| | | | dBtgh | | 20 | 0 | 1 | 0 | Not applicable | 2 | 17 |
| | | | n.i. | | 23 | 4 | 0 | 0 | Not applicable | 2 | 17 |
| E7 | 25 epi + 26 ExE | $10.5*10^6$ | Btgh | epi | 13 | 1 | 0 | 0 | 0 | 0 | 12 |
| | | | | ExE | 13 | 1 | 0 | 0 | 2 | 0 | 10 |
| | | | dBtgh | epi | 12 | 0 | 0 | 0 | 0 | 0 | 12 |
| | | | | ExE | 13 | 0 | 0 | 0 | 0 | 0 | 13 |

### Table S4. Primers used for PCR genotyping of E7 Btgh/dBtgh-injected embryos' cDNA

| **Gene target** | **Primer direction** | **Sequence (5'-3')** |
|---|---|---|
| Xist (cDNA) | forward | TCTATCTTGTGGGTCCTGGAG |
| | reverse | CTCCTCTAAATCCAGGCAATCC |

| | | |
|---|---|---|
| Ddx3y (cDNA) | forward | TGGAGGAGGAAATACAGAGAGC |
| | reverse | GGAGGACAATTATTTCCAGTTGC |
| Eif2s3y (cDNA) | forward | TGGCTGTGAAGTTGATGACC |
| | reverse | CCTTCTGTACGTACACCTAGG |
