## Supplemental Figures for "Perturbing nuclear glycosylation in the mouse preimplantation embryo slows down embryonic growth"

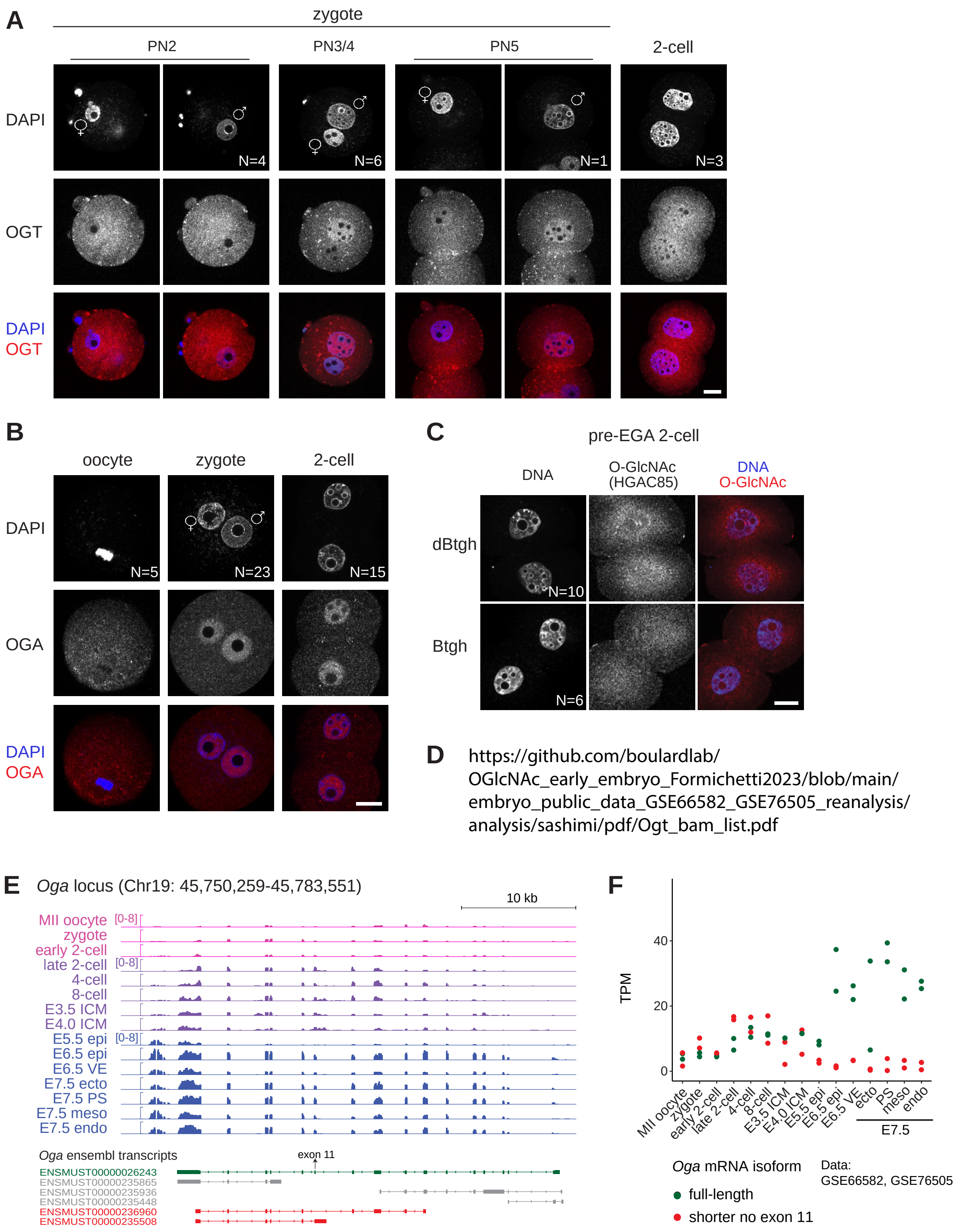


**Figure S1. Uncoupled dynamics of OGT and O-GlcNAc across mouse preimplantation development.** *legend on next page*

**Figure S1. Uncoupled dynamics of OGT and O-GlcNAc across mouse preimplantation development.**

(A) Immunofluorescence staining of OGT (ab177941) at different zygotic pronuclear stages and in 2-cell embryos collected after natural mating throughout the day of plug and 40 h post-hCG, respectively.

(B) Immunofluorescence staining of OGA in MII oocytes, zygotes and 2-cell embryos (22-26 h post-IVF) generated through IVF.

(C) Immunofluorescence staining of the O-GlcNAc modification in 2-cell embryos from zygotes injected with Btgh or dBtgh, using a different antibody than in Figure 1 (HGAC85) which shows signal at perinucleolar foci. Both effective O-GlcNAc removal and specificity of the O-GlcNAc signal at perinucleolar foci are confirmed by signal disappearance after Btgh injection, observed in all the 10 imaged Btgh-injected embryos.

(A-C) Embryos were mounted on coverslips and imaged using a scanning confocal microscope. Scale bar indicates 20 μm. PN = pronuclear stage. DNA was stained with DAPI. The total number of embryos imaged is indicated for each stage and experimental group and for (B,C) it comes from two independent IVF experiments. One z-plane is shown for each embryo, except for some zygotes for which two z-planes are shown.

(D) Sashimi plot of the annotation-independent analysis of *Ogt* mRNA exon junctions from the same datasets as in Figure 1D and 1E. Numbers indicate the raw number of reads spanning each junction. For each stage, biological replicates are overlapped, except for the late 2-cell for which one of the two replicates was discarded because of several high contaminant peaks. Only one alternative splicing event is found, precisely inside intron 4, and only during preimplantation development from the late 2-cell stage.

(E) Genome browser view of the *Oga* gene showing the alignment of mRNA-Seq reads from GSE66582 (Wu et al, 2016) (MII oocyte to E4.0 ICM) and GSE76505 (Zhang et al, 2017) (E5.5 to E7.5). Read counts are normalized to bins per million. One biological replicate per stage is shown.

(F) Salmon quantification of the *Oga* transcript isoforms from the same datasets as in (E). The full-length isoform is the one containing all exons (ENSMUST00000026243). TPM of isoforms ENSMUST00000235508 and ENSMUST00000236960 were averaged (shorter no exon 11). ENSMUST00000235865, ENSMUST00000235448 and ENSMUST00000235936 were excluded from the plot because of no evidence of their presence based on a manual inspection of the Integrative Genomics Viewer (IGV) tracks.

(E,F) epi = epiblast, ecto = ectoderm, PS = primitive streak, meso = mesoderm, endo = endoderm.

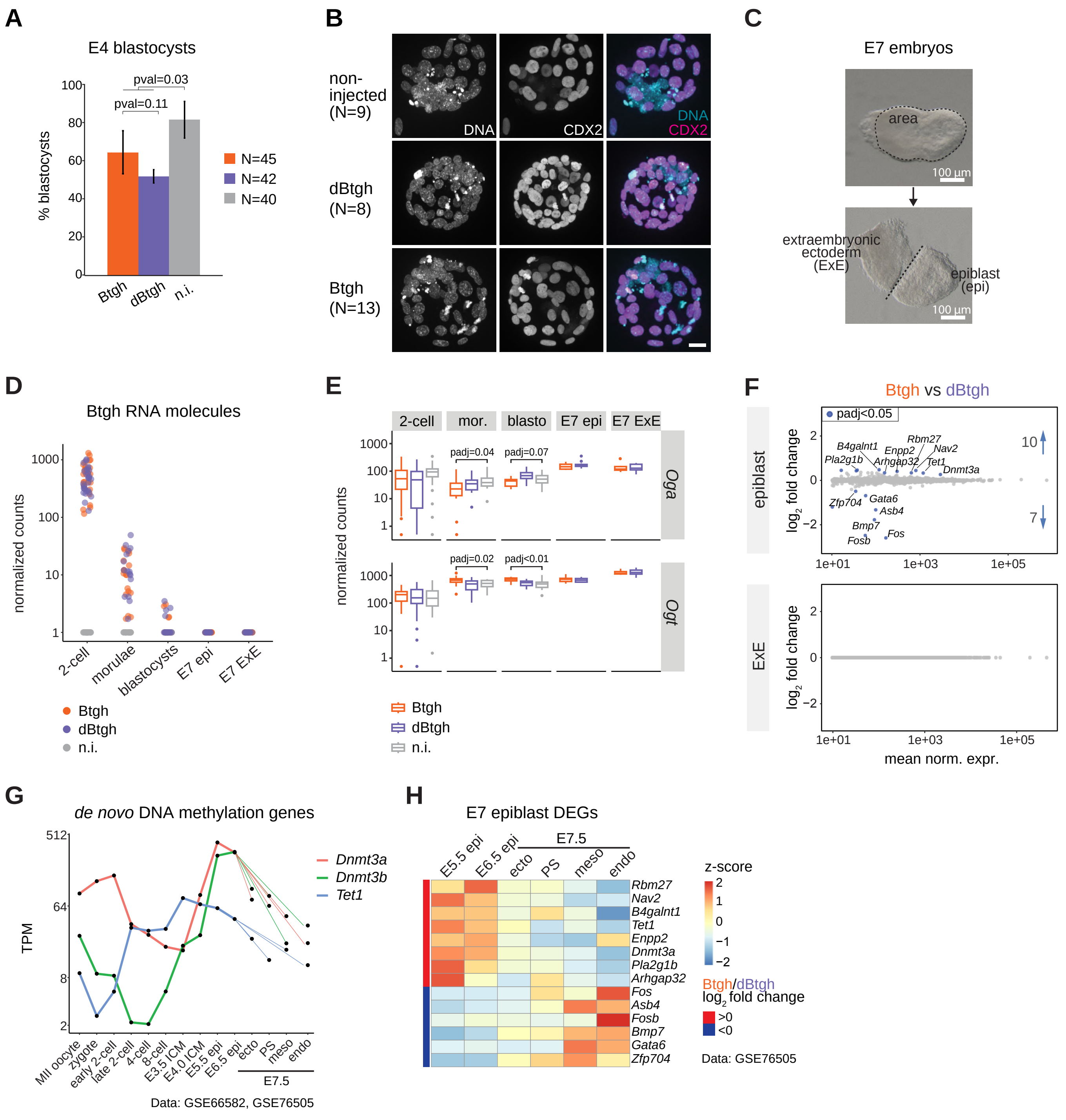


**Figure S2. Nuclear O-GlcNAc depletion does not affect differentiation but slows down development.** *legend on next page*

**Figure S2. Nuclear O-GlcNAc depletion does not affect differentiation but slows down development.**

(A) Percentage of healthy embryonic day 4 (E4) blastocysts which developed ex-vivo from Btgh/dBtgh injected and non-injected zygotes. Bar heights and error bars indicate the average and standard deviation, respectively, for four replicates of the microinjection experiment. The total number of starting 2-cell embryos is stated in the figure legend. The lower developmental rate of both injected groups is expected and due to the injection procedure. P-value was computed using unpaired Student's t-test, assuming unequal variance.

(B) Immunofluorescence staining of OGT and the trophectoderm marker CDX2 in blastocysts developed from Btgh/dBtgh-injected and non-injected embryos. Embryos were mounted in drops and imaged using a spinning disk microscope. Maximum projection of all z-planes is shown. Scale bar indicates 20 µm. DNA was stained with DAPI. The total number of imaged blastocysts is indicated.

(C) Trans-illumination image of a dissected E7 embryo. The dash line in the top image indicates how the area was measured as a proxy of the size of the embryo (shown in Figure 3C). The dash line in the bottom image shows the cut performed to separate the two halves, largely corresponding to the epiblast (epi) and extraembryonic ectoderm (ExE) tissues. The transcriptome of individual epi and ExE was analyzed using single-embryo mRNA-Seq.

(D) DESeq2-normalized counts of Btgh/dBtgh RNAs at all embryonic stages analyzed in this study. Y-axis ticks are in log10 scale.

(E) DESeq2-normalized counts of *Oga* and *Ogt* at all embryonic stages analyzed in this study, showing the compensatory downregulation of *Oga* and upregulation of *Ogt* in morulae and blastocysts upon depletion of nuclear O-GlcNAc. padj = adj. p-value computed using DESeq2 Wald test and corrected for multiple testing using the Benjamini and Hochberg method. Y-axis ticks are in log10 scale.

(F) MA-plots from DESeq2 differential expression analysis of E7 epiblasts or ExE from Btgh-injected embryos versus dBtgh-injected ones. Only genes with mean of DESeq2-normalized counts ≥ 10 are shown. All genes with adj. p-value < 0.05, any log2FC are colored and labeled (except three pseudogenes) and their number is indicated.

(G) Expression of *de novo* DNA methyltransferases *Dnmt3a* and *Dnmt3b* and DNA 5-methylcytosine hydroxylase *Tet1* throughout mouse embryonic development (mRNA-Seq data from GSE66582 (Wu et al, 2016) and GSE76505 (Zhang et al, 2017)). The two biological replicates per stage were averaged. Y-axis shows Transcripts Per Million (TPM) and ticks are in log2 scale.

(H) Heatmap of the expression of all differentially expressed genes (DEGs; adj. p-value < 0.05, any log2FC) in Btgh-injected epiblasts (labeled in (F)) at the transition between E5.5 and E7.5 (mRNA-Seq data from GSE76505 (Zhang et al, 2017)). TPM values were log2-transformed and scaled by rows.

(D,E,G,H) epi = epiblast, TE = trophectoderm, ecto = ectoderm, PS = primitive streak, meso = mesoderm, endo = endoderm.

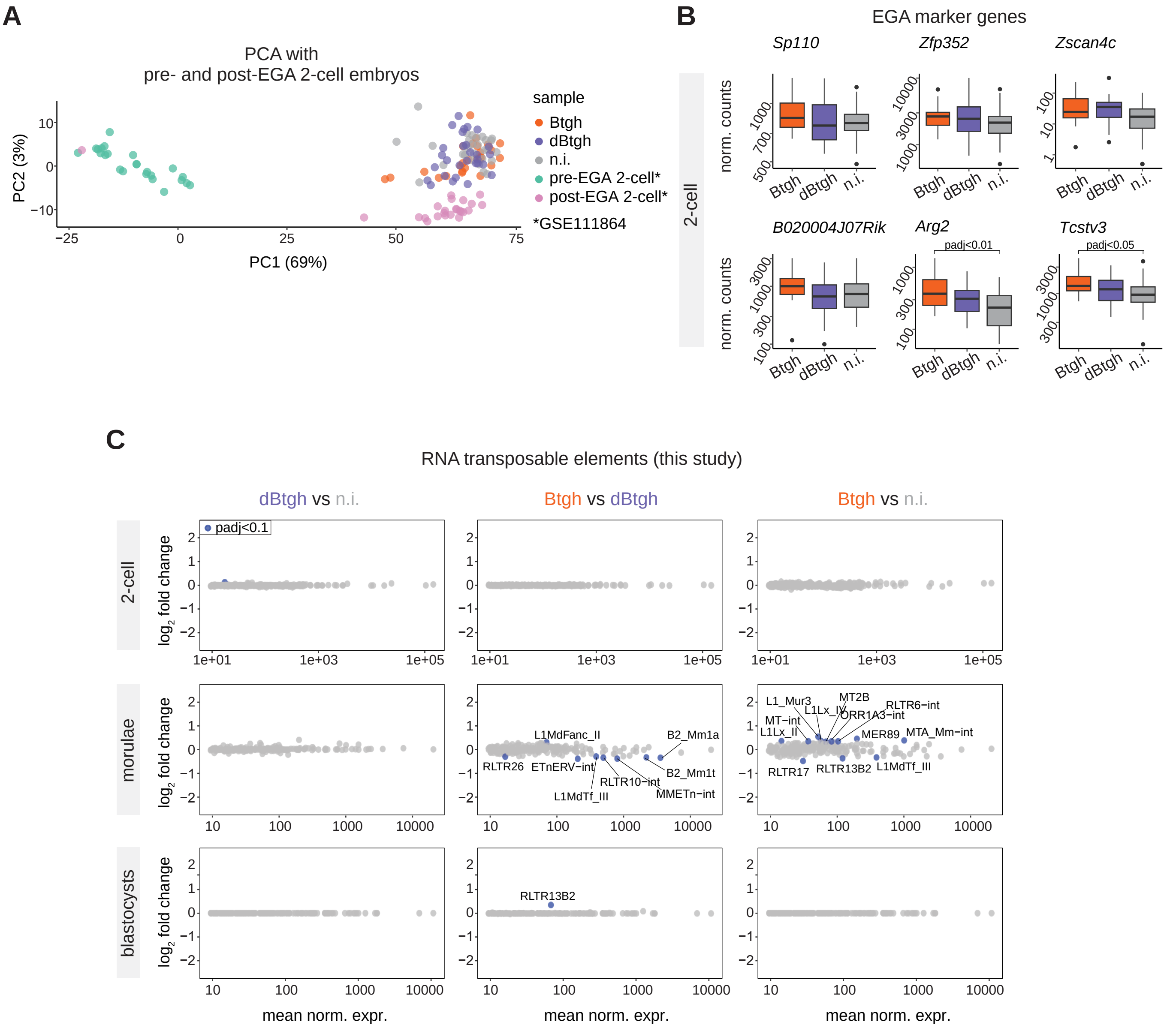


**Figure S3. Expression of retrotransposons in nuclear O-GlcNAc-depleted and unperturbed preimplantation embryos.**

(A) PCA of the three experimental groups of 2-cell embryos from this study, together with pre-EGA and post-EGA 2-cell embryos generated through ICSI (GSE111864) (Conine et al, 2018). The 200 EGA+maternal genes with the highest variance in the GSE111864 dataset were used to perform the PCA.

(B) DESeq2-normalized counts of six EGA-associated genes in the three indicated experimental groups of single 2-cell embryos. Padj = adj. p-value computed using DESeq2 Wald test and corrected for multiple testing using the Benjamini and Hochberg method. Y-axes ticks are in log10 scale.

(C) MA-plots from DESeq2 differential expression analysis of RNA transposable elements between the three experimental groups of embryos at the three preimplantation stages. Only retrotransposons with mean of DESeq2-normalized counts $\geq 10$ are shown. All retrotransposons with adj. p-value $< 0.1$, any log2FC are colored, the ones with absolute log2FC $\geq 0.2$ are labeled.

*(continued on next page)*

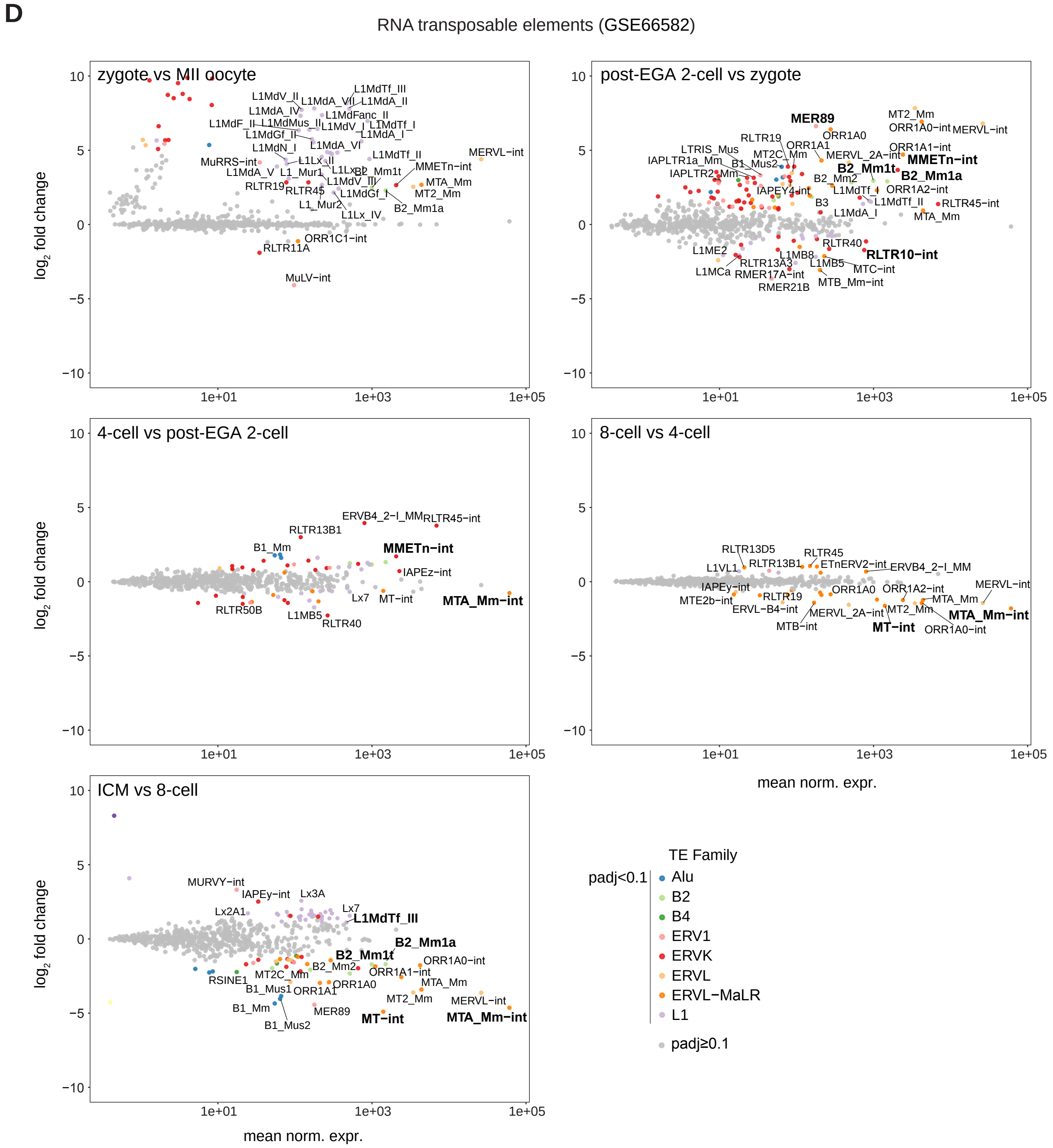


**Figure S3. Expression of retrotransposons in nuclear O-GlcNAc-depleted and unperturbed preimplantation embryos.** *(continued)*

(D) MA-plots from DESeq2 differential expression analysis of RNA transposable elements between successive stages of preimplantation development in publicly available mRNA-Seq data GSE66582 (Wu et al, 2016). Pre-EGA 2-cell embryos were discarded because one replicate out of two showed high DNA contamination based on DNA transposons FPKM values. Retrotransposons with adj. p-value < 0.1 are colored by family, labeled if standing out, in bold if significantly deregulated after O-GlcNAc depletion (B). Note that, with this analysis, elements which are more abundant in the genome (such as LINE elements) could be particularly sensitive to overestimated expression due to genomic DNA contamination of the RNA-Seq library.

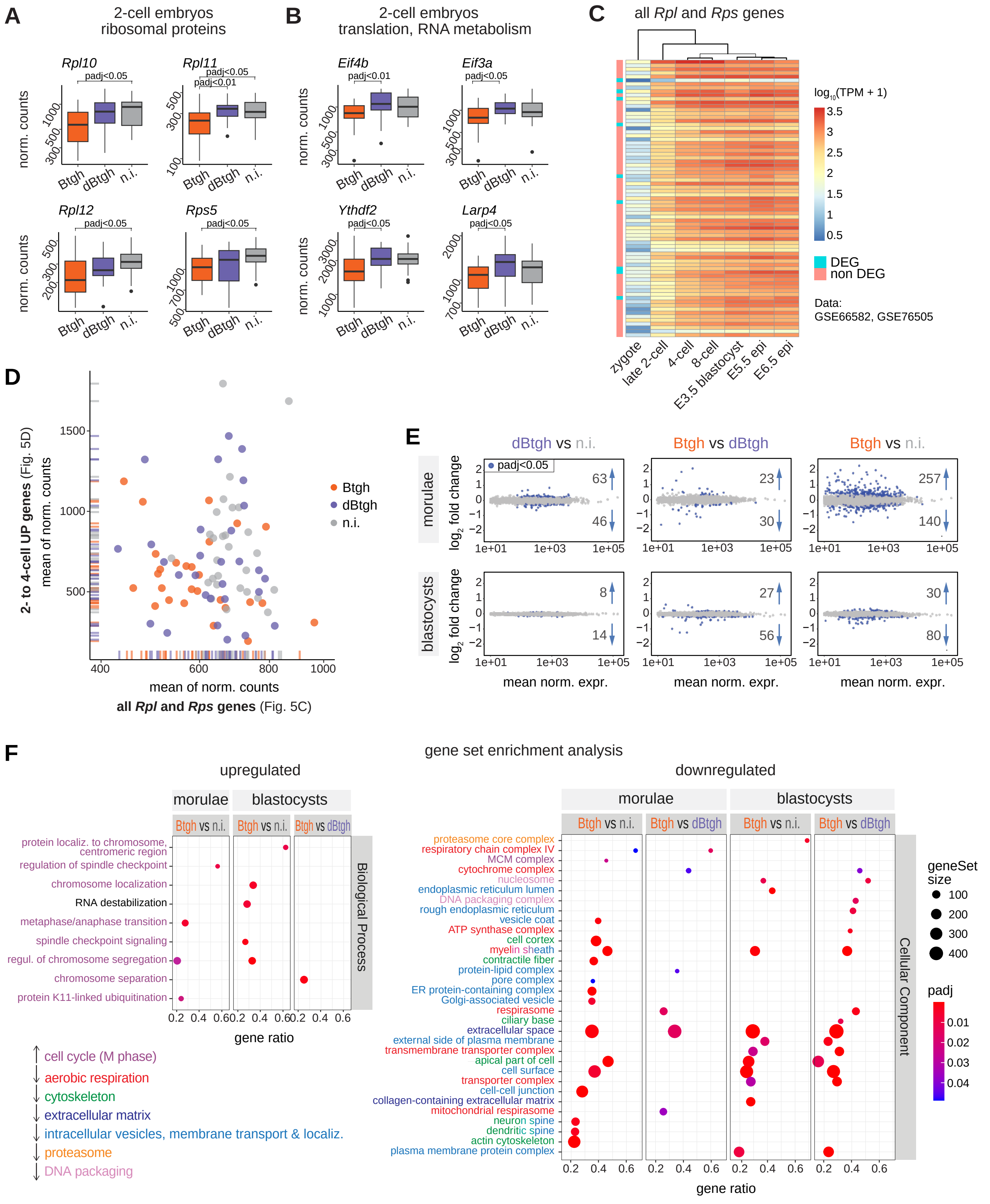


**Figure S4. Misregulation of mitotic and translation-related genes in nuclear O-GlcNAc-depleted embryos.** *legend on next page*

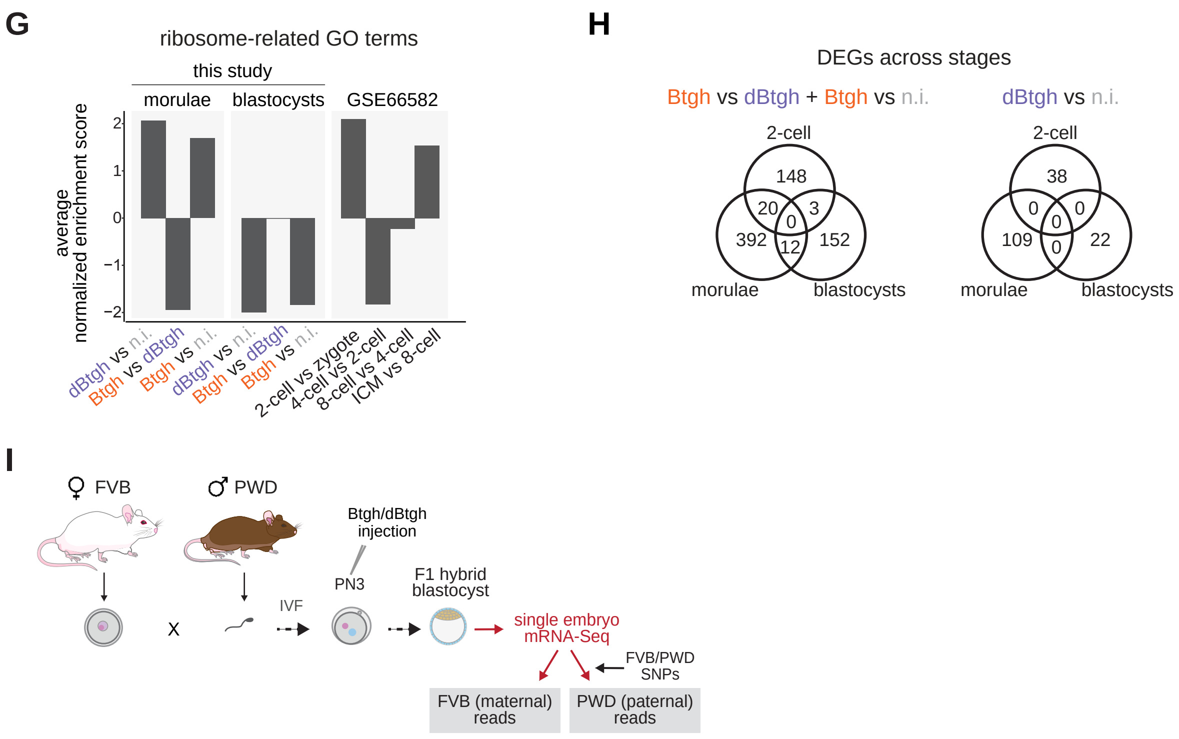


**Figure S4. Misregulation of mitotic and translation-related genes in nuclear O-GlcNAc-depleted embryos.**
(A,B) DESeq2-normalized counts of (A) four ribosomal proteins and (B) genes linked to RNA metabolism significantly downregulated in 2-cell embryos after nuclear O-GlcNAc depletion. padj = adj. p-value computed using DESeq2 Wald test and corrected for multiple testing using the Benjamini and Hochberg method.
(C) Heatmap of TPM values for all Mus musculus ribosomal protein genes from the Ribosomal Protein Gene Database (Nakao et al, 2004). Both rows and columns are clustered based on Spearman correlation. Genes which are differentially regulated at any stage after nuclear O-GlcNAc depletion (DEGs) are indicated.
(D) Average expression of all ribosomal protein genes (Figure 5C) versus average expression of genes upregulated between the 2- and 4-cell stage (Figure 5D) in single embryos from the three experimental groups. One outlier non-injected embryo was excluded.
(E) MA-plots from DESeq2 differential gene expression analysis between experimental groups of morulae and blastocysts. Only genes with mean of DESeq2-normalized counts $\geq 10$ are shown. All genes with adj. p-value $< 0.05$, any log2FC are colored, and their number is indicated.
(F) Gene set enrichment analysis of gene expression changes in nuclear O-GlcNAc-depleted morulae and blastocysts versus controls. Among the significant gene ontology (GO) terms, the upregulated Biological Process (BP) terms and the downregulated Cellular Component (CC) terms with the highest Normalized Enrichment Score are shown, ordered by gene ratio. Downregulated BPs and upregulated CCs can be found at
https://boulardlab.github.io/OGlcNAc_early_embryo_Formichetti2023/reports/Btgh_injected_blastocysts_SMARTSeq/GSEA_morulae_blasto_comparison.html.
Ribosome-related terms were excluded and plotted in (G). The size of dots is proportional to the number of total genes of a GO term. Gene ratio = fraction of total genes of the GO term which are concordantly changing between the two conditions. *(continued on next page)*

**Figure S4. Misregulation of mitotic and translation-related genes in nuclear O-GlcNAc-depleted embryos.** *(continued)*

(G) Average Normalized Enrichment Score for all ribosome- and translation-related terms found among significant terms in the GSEA results of (left) the morula and blastocyst datasets produced in this study and (right) a publicly available mRNA-Seq dataset spanning mouse preimplantation stages (GSE66582). 2-cell indicate post-EGA 2-cell embryos.

(H) Overlap among differentially expressed genes (DEGs; adj. p-value $< 0.05$, any log2FC) in Btgh-injected embryos versus any of the control groups (left) and in dBtgh-injected versus non-injected embryos (right) at the different preimplantation stages. DEGs were included only if DESeq2-normalized counts $\geq 10$ at the corresponding stage.

(I) Experimental scheme to produce F1 hybrid (FVB/PWD) embryos to assess whether depletion of nuclear O-GlcNAc affects paternal/maternal autosomal ratio as a measure of proper chromosome segregation.
